# The French flag problem revisited: Creating robust and tunable axial patterns without global signaling

**DOI:** 10.1101/2023.12.29.573602

**Authors:** Stephan Kremser, Gabriel Vercelli, Ulrich Gerland

## Abstract

Wolpert’s French flag problem conceptualizes the task of forming axial patterns with broad regions in multicellular systems. Wolpert described two different solutions to his problem, the balancing model and thresholding of a morphogen gradient, both of which require global, long-range signaling between cells. Since global signaling becomes challenging in large multicellular systems, we computationally explore alternative solutions, which use only local cell-cell signaling and are simple enough to potentially be implemented in natural or synthetic systems. We employ cellular automata rules to describe local signal processing logics, and search for rules capable of robust and tunable axial patterning with evolutionary algorithms. This yields large sets of successful rules, which however display only few types of behavior. We introduce a rule alignment and consensus procedure to identify patterning modules that are responsible for the different behaviors. With these modules as building blocks, we then construct local schemes for axial patterning, which function also in the presence of noise and growth, and for patterns with a larger number of different regions. The regulatory logic underlying these modules could therefore serve as the basis for the design of synthetic patterning systems, and as a conceptual framework for the interpretation of biological mechanisms.

## Introduction

One of the many remarkable features in the development of multicellular organisms is the formation of spatial gene expression patterns. The ability to spatially coordinate gene expression is a fundamental requirement for cell differentiation in animal and plant development, and spatial transcriptomics techniques now allow to measure the expression levels of genes systematically throughout tissue space, on a genome-wide scale (***Rao et al., 2021***). However, many of the principles and mechanisms underlying the formation of the spatial patterns still remain elusive. To explore such principles on a conceptual level, Lewis Wolpert introduced the ‘French Flag problem’ (***Wolpert, 1968***) as an abstraction of the task of forming axial patterns, i.e., quasi-one-dimensional patterns along a body axis. Specifically, the French Flag problem is to form a tri-partite pattern, such as that of a French flag, which is defined by the relative proportions of its different regions, rather than their absolute sizes (***Sharpe, 2019***).

In his seminal work (***Wolpert, 1968, 1969***), Wolpert proposed two different axial patterning mechanisms, conceptualized by the gradient model and the balancing model, respectively (***Sharpe, 2019***). The gradient model led to the notion of positional information encoded in the concentration gradient of a morphogen (***Wolpert, 1969***; ***Tkačik and Gregor, 2021***). In contrast, the balancing model represents a form of self-organized pattern formation, akin to the Turing mechanism (***Turing, 1952***) but without an intrinsic wavelength of the pattern (***Ishihara and Kaneko, 2006***). Both mechanisms have in common, however, that they require long-range signaling: In the case of the gradient model, cells respond to a globally diffusible molecule, for which a concentration gradient is maintained. In the balancing model, cells produce different long-range signaling molecules depending on their internal gene expression state, and cells adopt the state of their neighbors, if the total signal concentration emitted by them is weaker than their own.

Morphogen gradients and positional information are firmly and quantitatively established in model systems such as the early *Drosophila* embryo (***Tkačik and Gregor, 2021***). More generally, however, long-range signaling is limited by the diffusivity of the signaling molecule, such that alternative mechanisms are likely required for axial patterning over larger scales in multicellular systems (***Dickmann et al., 2022***). Alternative mechanisms are valuable also for the construction of synthetic multicellular systems ***Toda et al. (2018)***; ***Dupin and Simmel (2019)***. Mechanisms based on short-range signaling, between cells and their neighbors, are particularly promising in this context. On the one hand, short-range cell-to-cell communication, e.g., via Delta-Notch signaling (***Ehebauer et al., 2006***; ***Andersson et al., 2011***; ***Sjöqvist and Andersson, 2019***), Wnt signaling (***Logan and Nusse, 2004***), or Eph/Ephrin signaling (***Klein, 2012***), is ubiquitously available. For instance, the Notch pathway, which is conserved across the metazoa (***Bray, 2006***), uses membrane-bound proteins for communication between adjacent cells, regulating the formation of developmental patterns through lateral inhibition and lateral induction (***Bocci et al., 2020***). On the other hand, recent advances in synthetic biology enable the insertion of orthogonal short-range signaling systems into cells (***Toda et al., 2018***) and the creation of artificial cells with local communication (***Dupin and Simmel, 2019***).

Here, we revisit the French Flag problem to identify alternative mechanisms for axial patterning based only on short-range signaling. Towards this end, we use evolutionary searches to explore a large space of different models. These models have in common that they describe a one-dimensional array of locally communicating cells: Each cell is able to produce local signals, process signals from its neighbors, and respond in a context-dependent manner by switching its internal state. The models differ with respect to their local rules for signaling and signal processing. From a computational perspective, the French Flag problem with the short-range signaling constraint can be considered as a distributed computing problem — a number of cells jointly and distributively solves a task, without any of the cells receiving all of the inputs or observing all of the outputs (***Afek et al., 2011***). This computational perspective is reflected in our choice of the modeling framework for the signal processing of cells: We use the cellular automata (CA) framework, which was originally introduced and leveraged to study questions at the interface between the theories of computation and pattern formation (***Ulam et al., 1962***; ***Neumann et al., 1966***; ***Conway, 1970***; ***Wolfram, 1983, 1984***). Later, the CA framework also proved useful in describing the formation of cell differentiation patterns in various *in vivo* systems (***Akberdin et al., 2007***; ***Manukyan et al., 2017***; ***Adhyapok et al., 2020***).

By focusing on logical rules for local signaling and the dynamics of a small number of discrete cell states, CA provide a suitable level of description for our study. CA have previously been used to explore patterning principles for cellular systems (***Deutsch and Dormann, 2005***; ***Basanta et al., 2008***; ***Ramalho et al., 2021***; ***Bojer et al., 2022***), and to construct specific solutions for different “Flag problems” (***Herman and Liu, 1973***; ***Rohlf and Bornholdt, 2005, 2009***; ***Nichele et al., 2017***). Clearly, modeling frameworks other than CA are available, such as message passing models, which have been used to construct solutions to the French flag problem based on the communication and computation abilities of digital electronic devices (***Ancona et al., 2019***). As we will see below, the CA framework is well suited for an evolutionary optimization approach, since it naturally lends itself to a multiple alignment procedure that enables the identification of ‘consensus’ or prototype rules from an ensemble of selected CA models. This procedure proves instrumental to dissect successful patterning strategies into pattern formation modules, which each represent a relatively simple patterning principle that could conceivably be implemented in biological or bioengineered multicellular systems. We show that these modules serve as building blocks to engineer patterning schemes. Finally, we characterize the robustness of these schemes to growth of the system and noisiness of the systems’ dynamics or initial conditions, and show how strategies for different proportions of stripes can be either constructed or searched for.

## Model and Methods

### Modeling framework

We use cellular automata (CA) as conceptual models for tissues or synthetic systems composed of individual cellular units, which communicate with their neighbors via signaling mechanisms. This modeling framework focuses on the information processing within cells. It thereby enables our investigation of different *strategies* for pattern formation via contact-based signaling, regardless of the underlying molecular mechanisms. Computationally, the simplicity of CA allows the exploration of a large class of models, and evolutionary searches for models that satisfy different criteria.

Given our focus on axial patterns, it is sufficient to consider one-dimensional CA composed of a line of cells. Axial patterning in two-dimensional systems can use the same strategies as in one dimension, while an additional lateral coupling between adjacent lines of cells could increase the robustness of the patterning process (***Ramalho et al., 2021***). The cells in a one-dimensional CA of size *L* are labelled by a position index *i* = 1 … *L*. Each cell *i* has an internal state, represented by a single integer variable *x*_*i*_ ∈ {0, …, *k* − 1}, where *k* denotes the number of different internal states. For the French flag problem, the minimal number of states is *k* = 3, but we will also consider CA with more states. For instance, a fourth state could correspond to an initial, undifferentiated state, or to a transient state that does not occur in the final pattern.

In the CA modeling framework used here, cells do not change their relative position *i* in the system. However, cells can change their internal state in a context-dependent manner, i.e., depending on their current state and the current state of their neighbors. These state transitions occur at discrete time points and are governed by an update function *f*, called rule (fig. 1a). In a biological interpretation, the rule subsumes the effect of local inter-cellular interactions, intracellular signal processing, and internal differentiation processes. Computationally, the update rule takes as an input the states 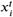 of a cell *i* and its nearest neighbors at a given time *t*, and outputs the new state of cell *i* at time *t* + 1,

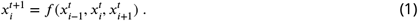

**Figure 1.**
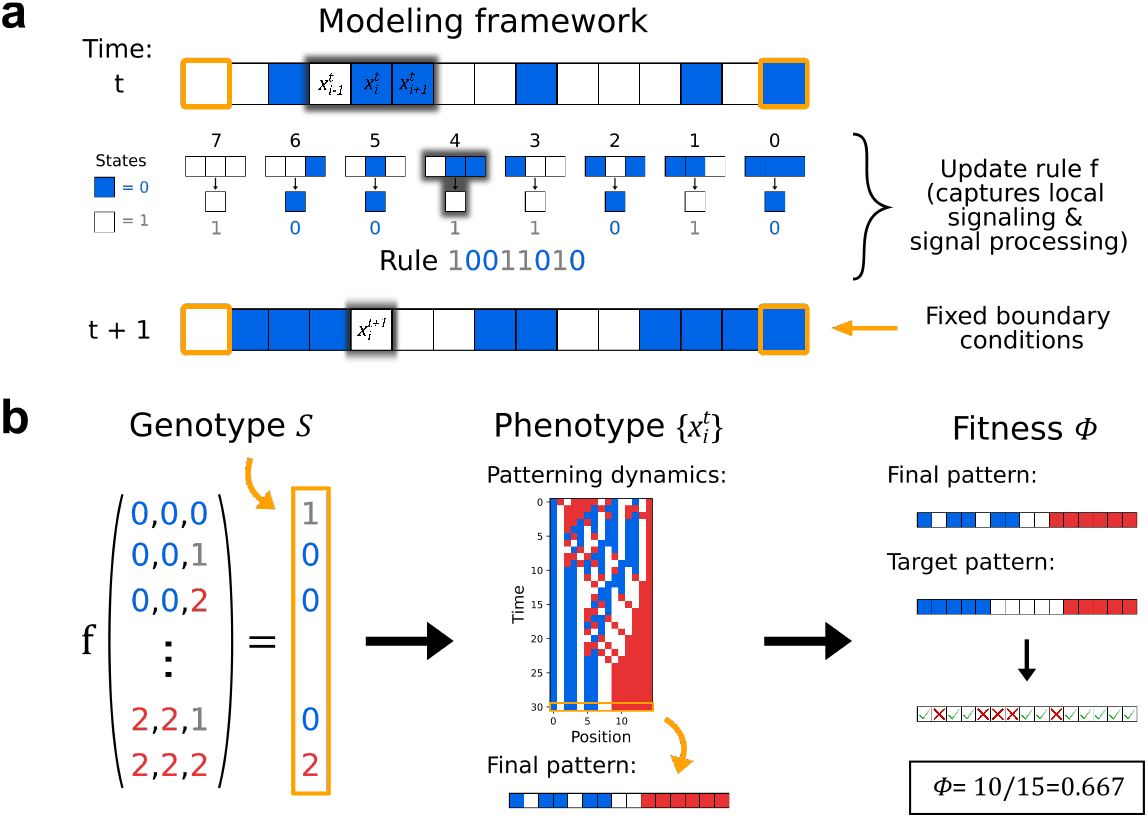
Cellular Automata (CA) as a minimal model of pattern formation via local cell-cell signaling. (a) Example of a one-dimensional CA of length *L* = 15, two states and fixed boundary conditions. Given a pattern of cells in the blue (0) or white (1) state at time *t*, an update function *f* (the “update rule”) determines the pattern at time *t* + 1. This rule maps each of the 8 possible configurations of a cell with its two nearest neighbors to an output, the new state of the cell. Each rule is uniquely identified by the string of output values for all possible input configurations in a prescribed order. (b) By analogy to pattern-forming developmental systems, we define a genotype to phenotype to fitness relation for the CA models. All cells in the system use the same update rule, which specifies the dynamical behaviour of the system. The string of output values encoding the rule is thus interpreted as a genotype, whereas the pattern formation process produced by the rule is the phenotype. While the fitness could generally depend on various aspects of the pattern formation process, we assume here that fitness depends only on the final pattern: The fitness is defined as the fraction of the final pattern that matches the target pattern. In the shown example, the French Flag pattern is the target pattern, but other axial patterns are also considered.

Each update rule *f* is uniquely specified by the sequence *S* of digits that represent the rule’s outputs to all possible combinations of input signals. This sequence also serves as the identifier for the rule (fig. 1a). Here, we only consider CA that apply the same rule to all cells, such that the behavior of a cell does not explicitly depend on its position *i* in space.

The update rule (1) is deterministic, since the current cell states 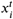 uniquely determine the cell states at subsequent times. In contrast, actual cells and synthetic cell-like systems are subject to biomolecular noise, which contributes a probabilistic component to their behavior. In our modeling framework, we explore the effects of noise using probabilistic updates of the form

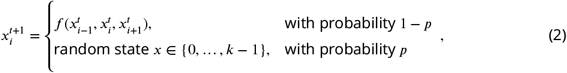

where the cell follows the context-dependent update rule with a (high) probability 1 −*p*, but ignores this rule with a (small) probability *p* by randomly choosing any state with equal likelihood.

While conventional CA models assume a fixed grid of cells and updates that happen simultaneously for all cells, biological systems often grow and their constituent cells are not perfectly synchronized. We therefore also analyze the effects of growth and asynchrony on pattern formation within our modeling framework. Towards this end, we allow for different levels of asynchrony by setting a probability *p*_*A*_ for a single cell to undergo an asynchronous update per time step. For each update, we determine, according to the probability *p*_*A*_, which cells to update asynchronously, and carry out the synchronous update for the remaining cells. We then perform the asynchronous updates in random order. The probability *p*_*A*_ then serves as a continuous parameter to control the degree of asynchrony. To introduce growth, we define an analogous parameter, the probability *p*_*divide*_ per time step for each cell to divide. The cell divisions that are selected according to this probability are carried out at each time step after the updates of the internal states, with daughter cells inheriting the internal state of their respective mother cells at that time.

Iterating the update procedure described above generates a timeline of state patterns, which is a characteristic of the pattern formation process under a given rule. From a biological perspective, the sequence *S* representing the update rule can be thought of as a genotype, which encodes a patterning strategy. The state *x*_*i*_ of a cell can be regarded as a cell type, and the timeline of state patterns 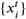 will be regarded as the phenotype of the respective rule (fig. 1b). Note that this phenotype is determined by both the rule and the initial pattern – the initial state 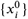 of all cells. So, in the biological analogy, the phenotype does not only depend on the genotype *S*, but also on the epigenetic information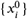.

While the phenotype is a pattern formation process, we are especially interested in the final pattern of cell states in the context of the generalized French flag problem considered here. Since in biological systems the time to develop a pattern is limited, we follow the dynamics of the cellular automaton for up to *T* = 2*L* steps in a system of size *L*, and compare this final pattern to the target pattern (alg. S1). The rule dynamics can, in principle, be transient or stationary, but we later show that our procedure produces rules that form stationary patterns with a time to steady state not larger than 2*L*.

### Fitness

To quantify how close the final pattern of the pattern formation process is to the target pattern, we use a fitness function. The closer the final pattern resembles the target pattern, the larger is the fitness Φ, which is defined as the percentage of positions at which the final pattern matches the target pattern (fig. 1b),

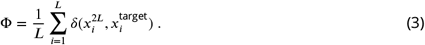

Because a genotype can produce different final patterns depending on the initial condition, we also consider for each genotype a mean fitness corresponding to the fitness average across possible initial conditions of a given length. Hence, the mean fitness for a given genotype *S* is only a function of the system size *L*. In practice, sampling all initial conditions for large *L* is computationally intractable, so in what follows we estimate the mean fitness from a random sample of *N*_*sample*_ initial conditions (alg. S2).

### Evolutionary algorithm

Evolutionary algorithms (EAs) mimic natural evolution to solve an optimization problem, by evolving a population of trial solutions to the problem via cycles of random mutation and selection. The successful application of EAs to CA models has previously been demonstrated [Mitchell, M., Hraber, P., Crutchfield, J., 1994. Evolving cellular automata to perform computations: Mechanisms and impediments. Physica D 75, 361391] (***Rohlf and Bornholdt, 2009***). Here, we use EAs to optimize the fitness Φ of pattern-forming CA rules. We start with a population of rules and use each of them to simulate CA with a fixed system size (default: *L* = 100), starting from *N*_*sample*_ different initial conditions (default: *N*_*sample*_ = 50). The random initial conditions are newly generated in each cycle of the EA and for each rule. Given the small sample size, this adds a stochastic element to the computation of the mean fitness according to eq. (3). For each rule, we apply two statistically independent mutations in its genotype, and determine its mean fitness as described above. If this mean fitness is higher than that of the original rule, we replace the original rule by its mutated version. Otherwise, we keep the original rule in the population. Finally, we sort the rules according to their fitness and discard the bottom 30%, replacing them by randomly generated rules (fig. 2a). This completes one evolution cycle and creates the initial population for the next one (see alg. S1).

**Figure 2.**
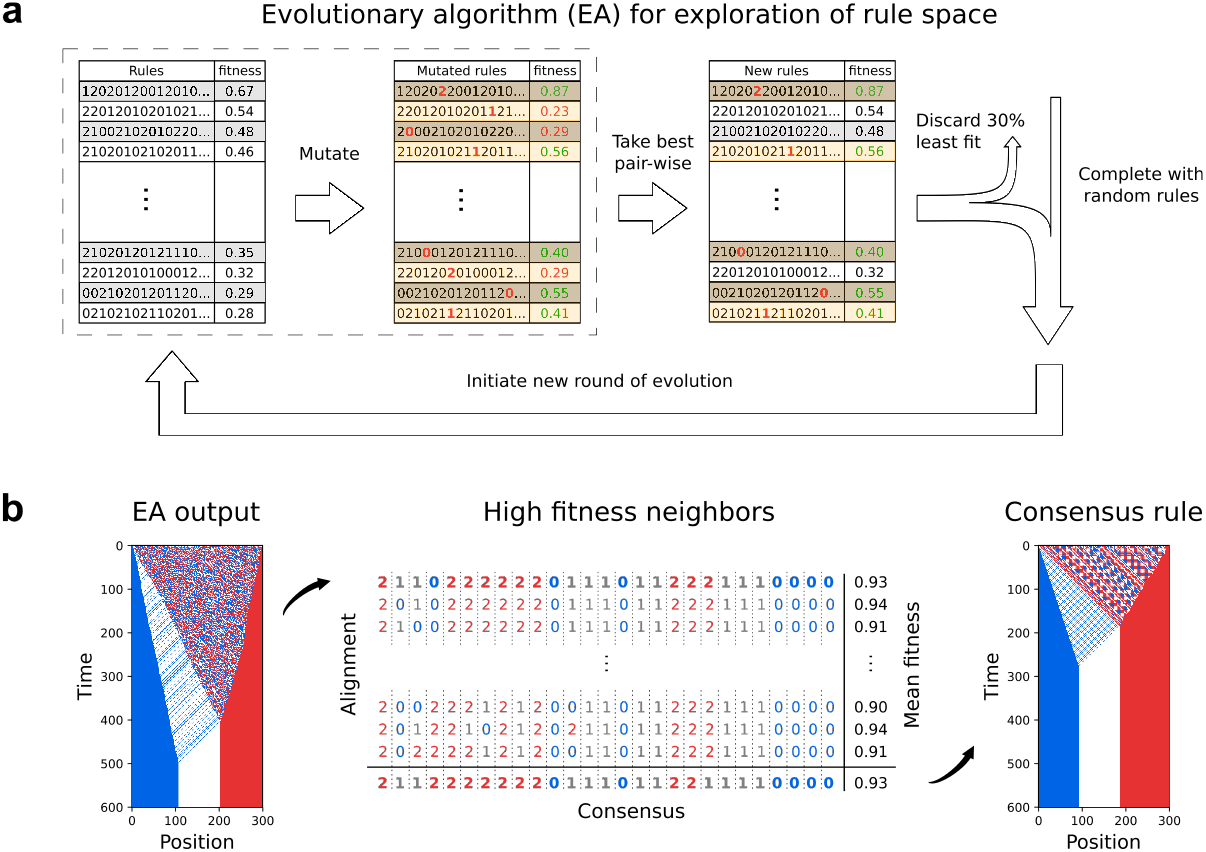
Evolutionary search for high-fitness rules and multiple alignment. (a) Schematics of the evolutionary algorithm (EA). Starting with a population of rules, the algorithm uses mutation and selection steps to find rules with higher fitness. (b) Left: Phenotype of a high-fitness rule found with the EA. Center: Multiple alignment of the genotype of this rule with those of its high fitness neighbors (Φ > 0.9) up to a Hamming distance of 6, totalling 379 different rules (the first rule in the alignment is the EA output). The consensus rule is the result of the multiple alignment. Right: Phenotype of the consensus rule.

### Adaptive fitness calculation

When searching the whole rule space (fig. 3e-g), we calculate rule fitness (eq. (3)) using alg. S2 with an adaptive assignment of parameter values to improve computational efficiency: First, we calculate rule fitness by sampling *N*_*sample*_ = 10 initial conditions with length *L* = 30. If the fitness exceeds 2/3, 40 additional initial conditions with the same length are sampled and all 50 samples are used to calculate that rule’s mean fitness. This reduces the computational cost for rules that are likely low scoring. Only rules with fitness above 0.8 are saved.

**Figure 3.**
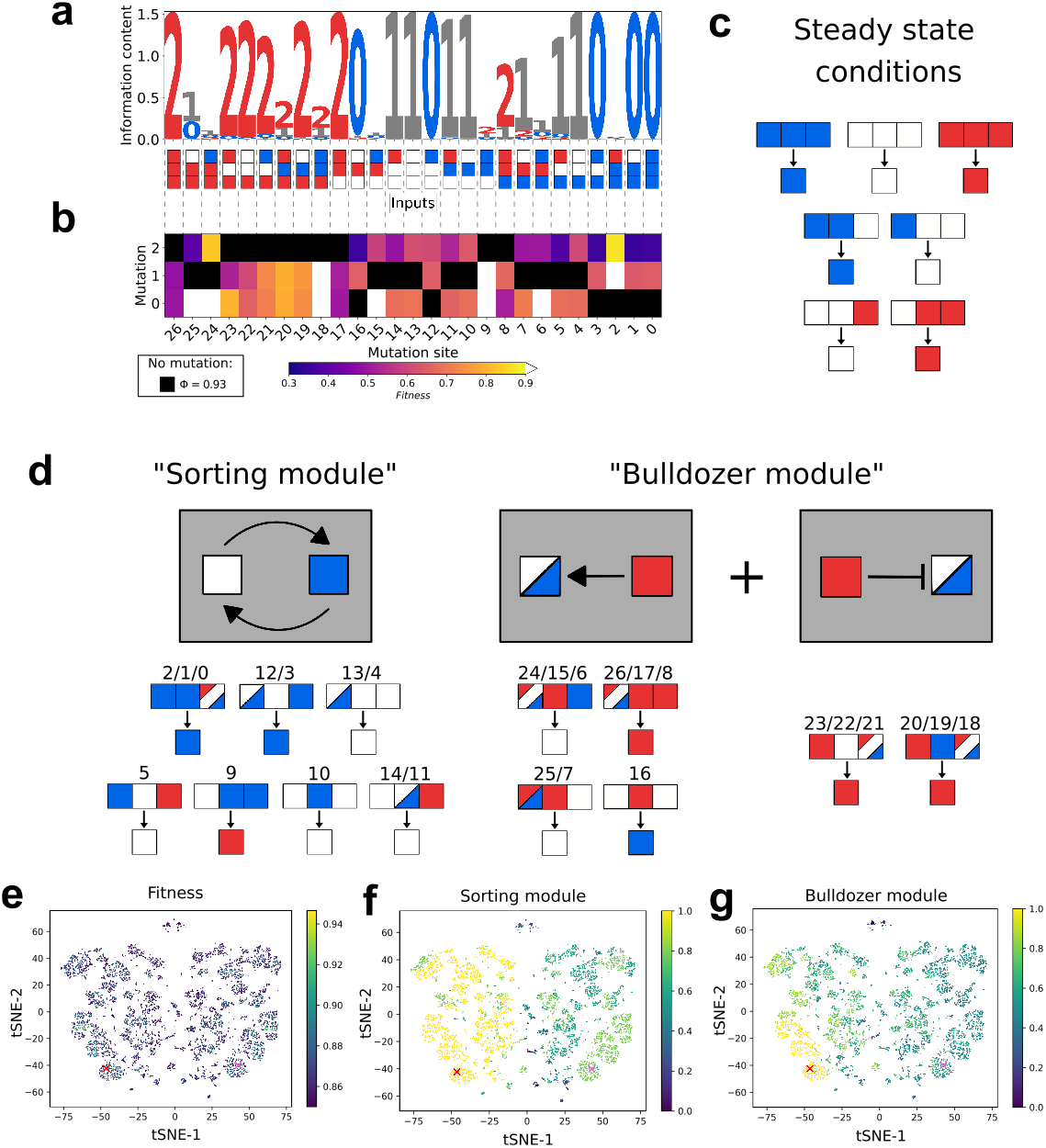
Dissecting the pattern formation mechanisms of the consensus rule. (a) Sequence logo for the consensus rule and its high-fitness neighbors (Φ > 0.9) up to a Hamming distance of 5 (in total 387 rules). The horizontal axis represents the 27 entries within the update rule (the input pattern is indicated for each entry to facilitate interpretation). The information content (decrease in Shannon entropy compared to a uniform distribution) in bits for each entry can be read off on the vertical axis as the total height of the stack of symbols. The relative height of the symbols represents their relative frequencies, with the most frequent output at the top. An entry of the update rule is considered as highly variable, if its entropy is more than 1 bit (information content less than *log*_2_(3) − 1 = 0.58). (b) Table of all single mutations of the consensus rule. Vertical axis shows to which state the output is mutated. The fitness of the mutant rule is color-coded (white indicates fitness above 0.9). Black squares mark the consensus output. (c) The condition that the target French flag pattern must be maintained once it is reached (steady-state) fixes seven entries of the update rule. (d) Partition of consensus rule entries into two patterning modules. The sorting module contains 12 entries and effectively implements a swap between white and blue states when white is to the left. The bulldozer module contains 15 entries and creates a red bulldozer state that moves to the right, while erasing any white and blue states to its right and alternately seeding blue and white states to its left. These two characteristics of the red bulldozer state are schematically represented by the grey boxes with the pointy arrow to white and blue states on the left, and the blunt-ended arrow to white and blue states on the right. (e-g) T-distributed Stochastic Neighbor Embedding (tSNE) plots of 3-state CA rules with fitness larger than 0.85, based on the genotypes *S* of the rules. Crosses represent the position of the consensus rule (red) and its symmetric partner (pink). The plots are colored based on rule fitness (e), overlap with conserved entries of the sorting module (f) or the bulldozer module (g). The left side of the plots in (f) and (g) shows high agreement with the sorting and bulldozer modules, while the right side shows high agreement with their respective symmetric partner rules (fig. S3c and d). Therefore, most high scoring rules combine the sorting and bulldozer module to achieve the French flag pattern.

## Results

### Evolutionary exploration of 3-state rules

We first apply the modeling framework described above to Wolfram’s classic French flag problem. The minimal number of states necessary to form a French flag pattern is *k* = 3. Even with only three states, the number of possible update rules is already vast: The three input cells of the rule, i.e. the central cell and its two neighbors (fig. 1), then each have three possible states, totaling 3^3^ different input configurations, each of which can be mapped to three possible output values, resulting in 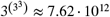 rules. To find French flag forming rules in a more efficient way than by exhaustive search, we used evolutionary algorithms (EAs) that apply iterative mutation, selection, and random insertion to a population of rules (see ‘Model and Methods’ and fig. 2a). The pattern formation starting from random initial conditions is scored by the fitness Φ defined by Eq. 3. The EAs lead to a gradual increase of the average fitness of the rules in the population, and a punctuated increase of the maximal fitness (fig. S1a,b). We found that including random insertions of new rules after selection slows the increase of the average fitness, but typically increases the maximal fitness that is reached (fig. S1b,c). We therefore settled on a single EA, in the following referred to as ‘Full EA’, which combines mutations with a fixed fraction of random insertions in each iteration.

The Full EA finds many rules in 3-state space, which have high fitness Φ and form final patterns resembling a French flag, starting from random initial conditions (fig. S1). Moreover, the evolutionary search is robust, in the sense that independent runs of the EA with different initial populations of rules yield rules with similar fitness (fig. S1c). The phenotype of the rule that displayed the highest fitness is shown in fig. S1d for three different system sizes. The fitness of this rule increases with the system size (fig. S1e), for reasons that will become clear below.

### Multiple alignment reveals a robust consensus rule

While the evolutionary search successfully identifies many high-fitness rules, understanding how the local signaling logic of those rules forms the target pattern remains a challenge. In order to better understand which properties of high-fitness rules are essential for their function, we use a multiple alignment procedure. This is conceptually similar to the identification of sequence motifs by aligning DNA, RNA, or protein sequences (***Bailey, 2008***), or network motifs by aligning biological networks (***Berg and Lässig, 2004***). In our case, we perform multiple alignments based on the genotypes *S* that define the update rules (fig. 2b). Each position in *S* encodes the rule’s output value for a specific configuration of the three input cells. When the multiple alignment is performed on related (“homologous”) high-fitness rules, a conserved value in a column of the multiple alignment thus indicates that the encoded context-dependent cell state transition is important for the patterning mechanism implemented by these rules. By contrast, columns in the alignment displaying highly variable values indicate input configurations that likely play non-essential roles. As for sequence motifs, we determine a “consensus rule” by selecting the most frequent value in each position of the multiple alignment.

To create an ensemble of related high-fitness rules, we take the genotype of one of the rules found by the evolutionary algorithm as a seed, and then select all rules that are at most 6 mutations away from this seed and have a fitness above a threshold of Φ = 0.9 (fig. 2b). This yields an ensemble of around 400 rules, with which we perform the multiple alignment to determine the consensus rule (fig. 2b). Note that this consensus rule is not the fittest rule in the ensemble of related high-fitness rules (fig. 2b). We will see further below that the consensus rule serves as a prototype, which exemplifies the general patterning strategy of the ensemble, but does not feature specific optimizations used by the highest-fitness rules. In the example shown in fig. 2b, the consensus rule has approximately the same fitness as the seed rule, but its phenotype 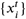 appears more ordered and reaches the steady-state more rapidly.

Visual inspection of the phenotypes 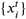 in fig. 2b also reveals that the seed rule and the consensus rule share a basic feature: The underlying patterning strategy is asymmetric, in the sense that the mechanism generating the red stripe is apparently different from the mechanism generating the blue stripe. We will dissect this patterning strategy in the next section, but emphasize already now that all rules within a multiple rule alignment must be from the same symmetry class in order for the consensus rule to be meaningful. For instance, it is straightforward to construct a symmetric partner of the consensus rule, for which the mechanisms to generate the blue and red stripes are interchanged (fig. S2). Even though this symmetric partner rule implements the same patterning strategy overall, its genotype *S* is not homologous to that of the consensus rule. This example illustrates why the multiple alignment cannot simply be constructed from all high-fitness rules identified by the evolutionary algorithm.

Our procedure to identify the consensus rule involves several parameter choices, including the fitness threshold that defines high-fitness rules, the choice of the starting rule to generate the ensemble, and the maximum Hamming distance from the starting rule. To test whether the consensus procedure is robust, we varied these choices and recalculated the consensus rule (table S1). The genotypes *S* of the obtained consensus rules displayed at most 5 mutations with respect to the consensus rule of fig. 2b. Out of all mutations, only one was within the conserved positions of the original multiple alignment, and this happened when the fitness threshold was decreased to 0.85, which includes many rules with inferior pattern formation. Taken together, these results indicate that the consensus procedure is not sensitive to the choice of the parameter values.

### Consensus rule can be dissected into two patterning modules

We now turn to the mechanistic interpretation of the consensus rule identified in fig. 2b. Towards this end, we first consider the sequence motif obtained from the multiple alignment. The sequence logo (fig. 3a) quantifies the sequence bias in the ensemble of related high-fitness rules, by depicting the statistical preferences at each site of the sequence in terms of the Shannon entropies. Out of the 27 sites in the sequence, 9 are perfectly conserved, 10 display a strong bias, and 8 are highly variable (fig. 3a). The pattern observed in the sequence logo is also consistent with the fitness effects of point mutations to the consensus rule (fig. 3b). Seven of the perfectly conserved sites in the motif can be understood to be a consequence of the hard requirement that the final French flag pattern is a steady-state pattern, i.e., it must be conserved under the update rule (fig. 3c).

A closer inspection of the consensus rule reveals that it can be partitioned into two functional modules (fig. 3d), which we refer to as the “sorting module” and the “bulldozer module”, respectively. Together, these two modules generate a French flag pattern, but they perform complementary functions. The sorting module effectively sorts blue states to the left and white states to the right. It contains all entries of the update rule in which the left and center positions of the input are not red (2 × 2 × 3 = 12 out of the 27 entries). Not all of these entries appear consistent with the simple sorting interpretation. In particular, entry #9 updates to a red state rather than white. However, this deviation is not essential (see below), which is indicated also by the fact that entry #9 is among the highly variable positions in the sequence logo (fig. 3a).

The bulldozer module contains the remaining 15 entries of the update rule (fig. 3d). Effectively, this module moves red states to the right until they hit another red state. In the process, a moving red state erases the pattern in front of it and generates new blue and white states in its trail (we therefore refer to the red state as ‘bulldozer state’). Through the repeated action of the bulldozer module, the red states accumulate on the right, forming the red stripe, while the last bulldozer leaves blue and white states behind, which are sorted by the sorting module to form the blue and white stripes in the left and center, completing the French flag pattern. This interpretation of the pattern formation process generated by the consensus rule suggests the following properties: (i) the total number of red states should be conserved during the process, (ii) the leftmost red state should seed blue and white states in a one-to-one ratio, (iii) the blue and white states to the left of the leftmost red state should be sorted into stripes maintaining their numbers, and (iv) the final pattern should scale with the system size, as required in the statement of the French flag problem. We checked these properties empirically (Appendix 4 and fig. S2). Property (iv) holds for the consensus rule, in the sense that the mean boundary positions scale linearly with the system size (fig. S2d). Properties (i) and (iii) become true when entry #9 of the consensus rule is mutated to white or blue (fig. S2a, b, g, h), because this stops the creation of additional red states. Property (ii) becomes true when entry #15 is mutated to blue (Appendix 4 and fig. S2c).

We created an analytically solvable model of the pattern formation mechanism with properties (i)-(iii). Using this model, we proved that the mean fitness of this mechanism converges to 1 as the system size *L* increases, with a power law exponent of −1/2 (Appendix 5). This also implies the scaling property (iv). Because the properties (i)-(iii) do not exactly hold for the consensus rule, it is not surprising that the mean fitness of the consensus rule converges to a value less than one (17/18) as the system length is increased (fig. S2f). However, the mean fitness of the rule where entry #9 is mutated to white does follow the predicted fitness scaling (fig. S2f).

Using the above insight, we can also rationalize the highly variable sites in the sequence logo (fig. 3a). These correspond to the entries controlling the bulldozer seeding process (#6 and #15), the dynamics of blue and white states in between bulldozers (#20, #24, #25), and inputs that rarely appear in the patterning process (#2, #9, #18). These entries are flexible not because they are completely irrelevant for pattern formation, but rather because there are different combinations of their outputs that successfully form a French flag. As an example, when entry #25 maps to white, most states in between red states are going to be white during pattern formation. This makes entries #6, #15, and #24, which require a blue state to the right of a red state, mostly irrelevant. Conversely, when entry #25 maps to blue, blue and white states appear often between red states, so inputs #6, #15, and #24 become relevant.

### Exhaustive search within 3-state rule space

Since the evolutionary exploration of the rule space used above may have missed other patterning strategies, we also sought to perform an exhaustive search of the whole rule space, trying to identify all successful patterning strategies. Such an exhaustive search is computationally challenging, but for the minimal case of *k* = 3 states it is feasible, in principle, since the number of rules is on the order of a trillion. In practice, this requires a very fast computation of the fitness of each rule. Towards this end, we used an adaptive algorithm (see Model and Methods), which quickly eliminates rules that clearly do not form a French flag and evaluates with more attention the remaining ones.

We used this exhaustive search approach to identify all high-fitness rules, defined here as rules with fitness estimates larger than *f* > 0.85 (in total 6675 rules). In order to assess and visualize to what extent these rules rely on the same types of “Sorting” and “Bulldozer” modules as the consensus rule, we chose the following strategy: We first represented the rules as dots in a two-dimensional plane, arranged such that similar rules lie close together. For this purpose, we used a t-distributed Stochastic Neighbor Embedding (tSNE) plot (***van der Maaten and Hinton, 2008***), based on the genotypes *S* of the rules (see fig. 3e). In this plot, we also marked the positions of the consensus rule (red cross) and its symmetric partner rule (pink cross). We then colored the dots within this same plot in different ways, according to color codes defined by different observables that characterize the behavior or structure of the rule. Note that the tSNE plot displays an approximate mirror symmetry, where nearly every cluster of dots on the left side has a symmetric partner on the right side (fig. 3e). The consensus rule and its symmetric partner rule are also in mirror symmetric positions on the left and right, respectively. This symmetry becomes useful when we next interpret the color patterns obtained from different observables.

Using fitness as the color code reveals no particular pattern in the tSNE plot (fig. 3e). However, when the dot color encodes the overlap of the rule with the blue-white sorting module (fig. 3f) of the consensus rule, the vast majority of rules on the left-hand side display full overlap. Here, overlap is defined as the fraction of module entries within the update rule that match the corresponding consensus module entry, not including the entries that were highly variable in fig. 3a (entries with information content less than log_2_ (3) − 1 = 0.58 bits). Many rules on the left-hand side also contain the red bulldozer module of the consensus rule, or have a high degree of overlap with it (fig. 3g). Conversely, the rules on the right-hand side display a high degree of overlap with the white-red sorting module and the blue bulldozer module of the symmetric partner rule (fig. S3). Color-coding also for the overlap with the steady-state conditions of fig. 3c reveals that a small group of rules at the top, as well as a few interspersed rules elsewhere, violate these conditions (fig. S3). Consistently, the average time to reach the steady state cannot be defined for these rules, and is shown as zero in the corresponding dot plot (fig. S3). These rules, which generate propagating wave fronts, received a high fitness score only because their transient pattern is typically close to the French Flag pattern at the fixed time point when the fitness is evaluated.

Taken together, our exhaustive search within 3-state rule space revealed no new patterning strategy beyond the strategy of the consensus rule, which combines a sorting module with a bulldozer module. In particular, the search revealed no “pure strategies”, based either only on sorting or only on a erase-and-reconstruct mode of patterning. We will next see that such pure strategies become possible when the rule space is expanded to four states.

### Pure sorting and landscaping strategies for axial patterning

Three is clearly the minimal number of cell states required to generate a French flag pattern. A fourth state, which occurs transiently but not in the final pattern, may offer the flexibility required to implement alternative patterning strategies. For instance, three states are insufficient to implement a pure sorting strategy, which starts from a random distribution of states and effectively moves all blue states to the left and red states to the right: The underlying issue is that local configurations of three neighboring cells in an inverted French flag state lead to an ambiguous update according to the sorting logic (fig. 4a). This issue can be resolved with the help of an additional transient state (shown in black), which mimicks the superposition of a blue and a red state. A resolution module involving this additional black state (fig. 4b) combined with three sorting modules (blue-white, white-red, blue-red) then successfully sorts an initial pattern containing the problematic inverted French flag configuration (fig. 4c,d). We refer to this 4-state rule encompassing the resolution module and the three sorting modules as the ‘Bubble rule’ (fig. S1a), because its behavior resembles that of the bubble sorting algorithm (***Knuth, 1998***), see fig. 5 (left).

**Figure 4.**
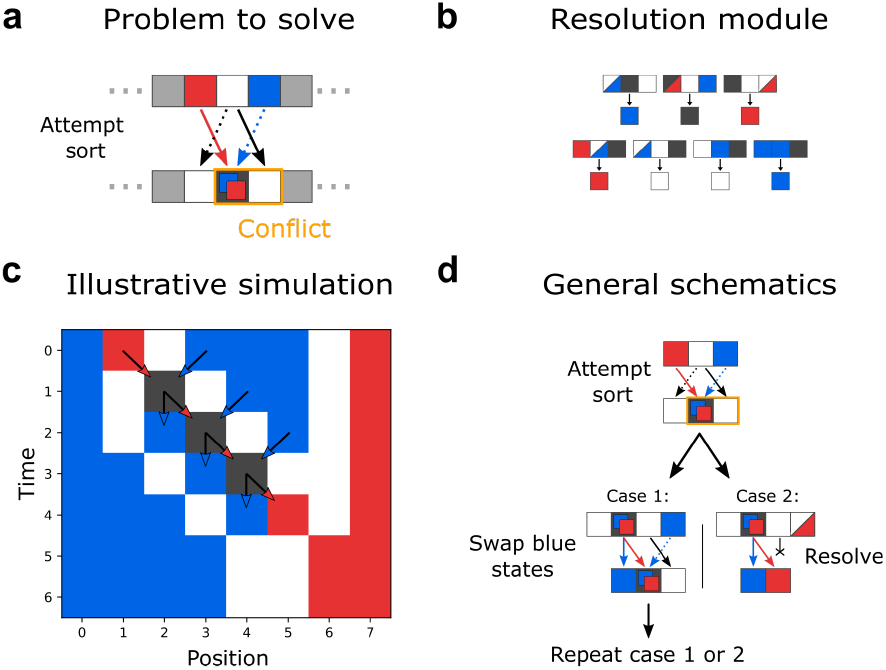
Pure sorting strategy. (a) Pure sorting with only three states faces a dilemma at local configurations of three cells in an inverted French flag pattern. According to the sorting logic, both lateral states, blue and red, should move to the center, while the white state should move both to the right and left. (b) The dilemma is resolved with an extra state (colored black, numbered 3 in genotypes *S*) representing the superposition of blue and red. A resolution module is formed by the shown set of updates. (c) Example of how the resolution module operates. A random initial pattern with local inverted French flag configuration is sorted, maintaining the initial number of cells in each state. To undo the duplication of the white state in the first time step, the resolution module lets the black state chase the extra white state on its right and, when they meet, exchange both states for a blue and a red state, resolving the ambiguity of the black state and erasing the duplicated white state. The arrows link this example to the general scheme underlying the resolution module shown in (d). An ambiguous position is always converted into a black cell and a duplicated white state. If the state of the cell to the right of the duplicated white is blue, that blue state is swapped with the one inside the black state (case 1). Otherwise, the red state takes the place of the duplicated white, resolving both the ambiguity of the black state and the duplication of the white state (case 2).

**Figure 5.**
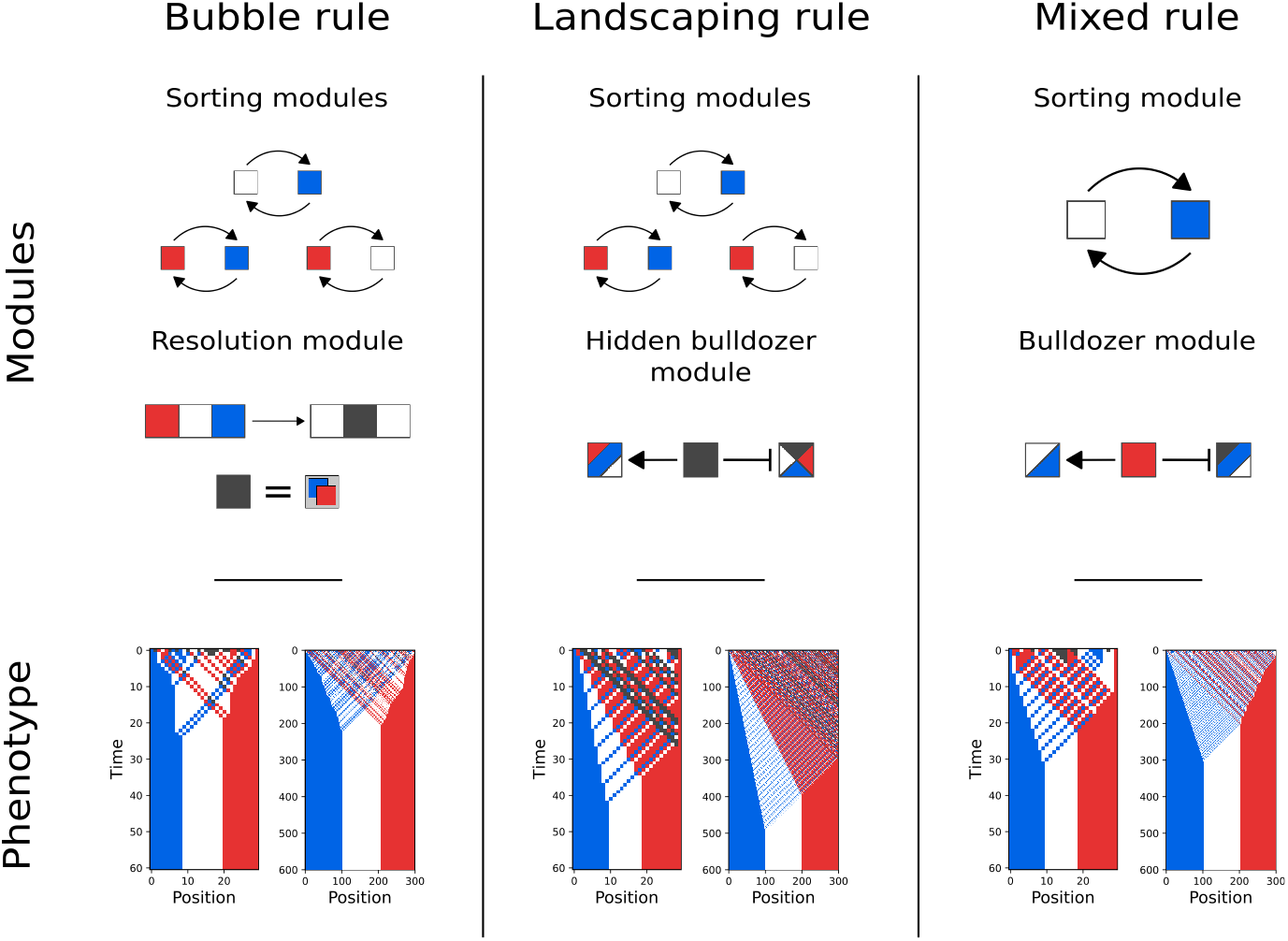
Three different axial patterning strategies in 4-state rule space. In addition to the mixed patterning strategy of the 3-state consensus rule, the introduction of a fourth, transient state also enables the formation of a French flag pattern via a pure sorting strategy (left), or via a full erase-and-reconstruct patterning strategy (middle). These new patterning strategies are implemented by combining the three pairwise sorting modules for the French flag color states with the resolution module or the hidden bulldozer module, respectively. The resulting phenotypic patterning behavior (bottom) is shown for two different system sizes, starting from random initial conditions in 4-state space. For comparison, a 4-state version of the 3-state consensus rule implementing the mixed patterning strategy is also shown (right). See fig. S1 for the full definitions of the three shown rules.

A full erase-and-reconstruct mode of forming the French flag pattern is also not possible with only three states. This is because the state that erases the initial pattern then has to be one of the states in the final pattern, such that it cannot erase itself. However, four states enable a “hidden bulldozer module” using a transient bulldozer state, which not only erases other states to its right, but also itself (fig. 5, middle). Compared to the bulldozer module of fig. 3d, this module has a modified seeding process, such that blue, white, and red states are seeded alternately to the left of the bulldozers. The states seeded by the bulldozer are then sorted into position by sorting modules for each pair of states. The problematic inverted French flag configuration never occurs, because of the seeding order of the bulldozer state. We refer to this 4-state rule combining the three sorting modules with a hidden bulldozer module as the “Landscaping rule”, as it re-patterns the entire system (fig. 5, middle).

The 4-state rule space also allows for a mixed patterning strategy akin to that of the 3-state consensus rule (fig. 5, right). Like the consensus rule, the 4-state “Mixed rule” (fig. 5, right) uses only a single sorting module for blue and white states and a bulldozer module with the red state as its bulldozer state. This 4-state bulldozer also erases the additional black state (see fig. S1 for the full rule definition). Taken together, we now have three different axial patterning strategies, which triggers the question how they differ with respect to properties such as the accuracy, speed, robustness, and tunability of the patterning process. We analyze each of these aspects in the following.

### Accuracy and speed of the patterning process

Our fitness function (eq. (3)) quantifies the accuracy of the patterning process by measuring the deviation of the final pattern from the target pattern. We therefore compare the accuracy of different patterning strategies by comparing their mean fitness values achieved across random initial conditions. A large mean fitness is associated with an update rule that forms the French flag pattern with high accuracy for most initial conditions. Figure 6 shows that the mean fitness grows with the system size *L*, for all three axial patterning strategies, approaching the maximal value 1 as a power law, supporting that all rules form perfect French flag patterns in the large system size limit. Interestingly, the Landscaping rule approaches this limit much faster than the other two, with a power law exponent around −1, compared to exponents close to −0.5 for the Bubble rule and the Mixed rule (inset).

**Figure 6.**
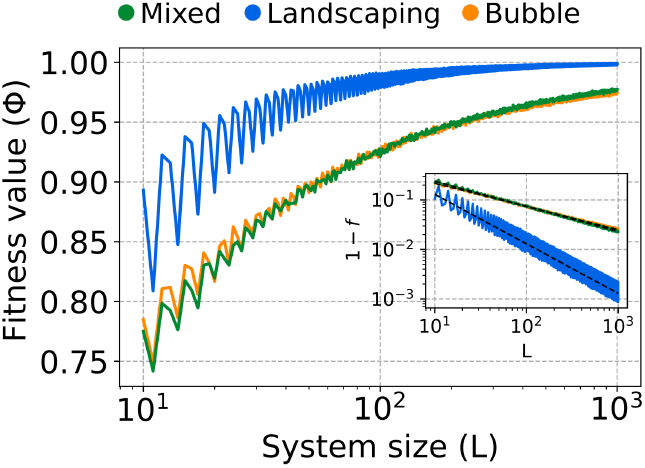
Comparison of the patterning accuracy of the three patterning strategies of Fig. 5. Accuracy is measured via the mean fitness value (averaged over 1000 randomly chosen initial conditions). The maximal fitness of one is achieved only when a perfect French flag pattern is formed for all initial conditions. The mean fitness increases with the system size, *L*, for all patterning strategies, but it approaches the maximal fitness with different power laws (inset). Linear fits on a log-log scale (inset) yield the exponents −.509 ± .001 for the Mixed rule, −1.00 ± .01 for the Landscaping rule, and −.469 ± .001 for the Bubble rule. The mean fitness oscillates with a period of three, since perfect French flag patterns can be formed only when *L* is divisible by three.

To understand the mechanisms underlying these behaviors, we used simple analytically solvable models to derive the fitness scaling of the rules from basic probability principles (Appendix 5). In short, the Bubble rule generates the final pattern by sorting initial cell states, such that the statistical fluctuations in the initial distribution of cell states propagates to the final state. These fluctuations scale as the inverse square root of the system size, explaining the exponent of the Bubble rule. The Mixed rule is similarly affected by the statistical fluctuations in the initial distribution of cell states, and thus displays the same scaling as the Bubble rule (see Appendix 5). In contrast, the Landscaping rule uses a bulldozer state to re-pattern the entire system. The fitness deficit is caused by the small number of states that escape this re-patterning because they appear to the left of the first bulldozer state. This is a constant number of states, on average, so they create a fitness deficit inversely proportional to the system size, explaining the power law exponent of the Landscaping rule.

Another important property of a patterning strategy is the time required to complete the pattern. To compare the patterning speed of the three 4-state rules of fig. 5, we analyzed the mean time to reach the steady state pattern, *T*_*ss*_(*L*), as a function of the system size *L*. We estimated *T*_*ss*_(*L*) from simulations, by averaging over different random initial conditions at a fixed *L*. For all three rules, *T*_*ss*_(*L*) scales linearly with *L*, however the rules display different linear scaling coefficients, i.e., different mean patterning speeds. The Bubble rule is fastest, with *T*_*ss*_ ≈ 0.74*L*, and the Landscaping rule is slowest, with a linear scaling coefficient of 1.67, while the Mixed rule is in between, with a coefficient of one (fig. S1). These scaling behaviors are also rationalized by the analytical models described above (Appendix 5). In a nutshell, the Mixed and Landscaping rules are slower, because they erase parts of the existing pattern before creating the target pattern, while the Bubble rule starts patterning immediately.

Taken together, the increased patterning accuracy of the Landscaping rule comes at the price of a reduced patterning speed. We will next see, in our analysis of robustness and tunability, that this is not the only trade-off between the different patterning strategies.

### Robustness of the patterning process

#### Robustness against noise-induced errors

In biological systems, noise is ubiquitous (***Tsimring, 2014***) and can be disruptive as well as constructive. More specifically in development, pattern formation can be affected by noise-induced errors in intercellular signal transmission, intracellular information processing, and cellular differentiation. Within the CA modeling framework, we describe such local errors by probabilistic updates, as defined in Eq. (2), where the error probability *p* serves as a measure of the noise level.

The three rules of fig. 5 respond differently to update errors (fig. 7a,b). When the perfect French flag pattern is used as the initial state, the pattern quickly disintegrates already at small error probabilities *p* under the Landscaping rule (fig. 7a). The Mixed rule is similarly sensitive to update errors, except for the red stripe, which is maintained better than the blue and white stripes. The Bubble rule is most robust overall to noise-induced update errors. These qualitative observations (fig. 7a) are consistent with a quantitative analysis of the error-induced shift in the boundary positions within the French Flag pattern (fig. 7b), which suggests that boundaries created by sorting modules are significantly more robust against update errors than boundaries created by bulldozer modules. The robustness of sorting modules is rationalized by noting that (i) any update error is sorted into the correct stripe, and (ii) the stripe width does not change on average, since a stripe of width *l* widens with a rate proportional to *L* − *l* and decreases width with a rate proportional to 2*l*, resulting in equilibrium lengths of *l* = *L*/3, generalizing a mechanism previously described for a two color pattern (***Rohlf and Bornholdt, 2005***). In contrast, bulldozer modules are sensitive to update errors, because erroneously generated bulldozer states re-pattern the system.

**Figure 7.**
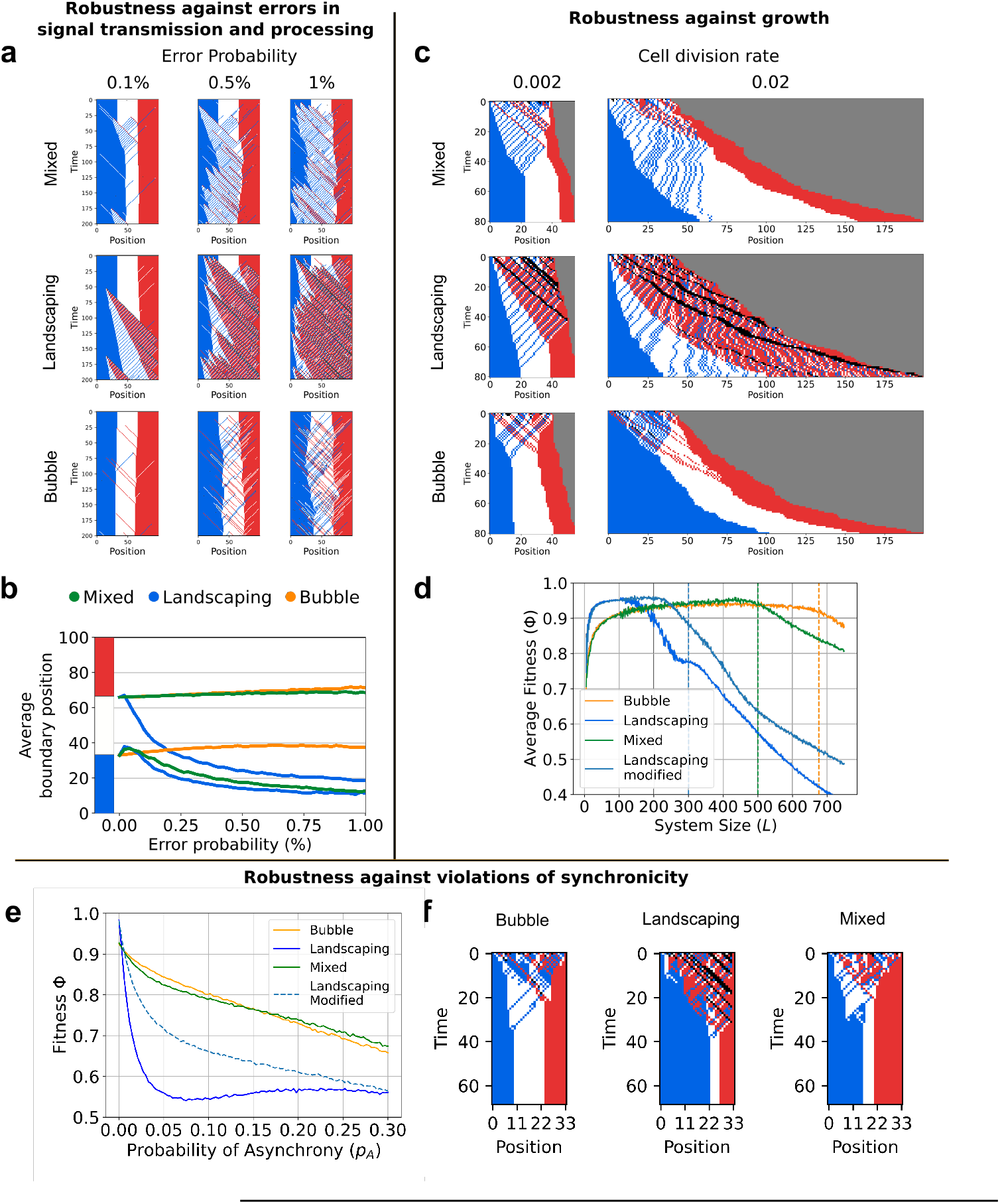
Robustness of the three patterning strategies of Fig. 5. (a) Maintenance of the French Flag pattern under noisy updates according to the Mixed, Landscaping, and Bubble rule. (b) Estimated average position of the two boundaries within the tripartite pattern, as a function of the update error probability *p*, for each of the three rules. Each point was calculated as an average over 1000 iterations starting from the French flag pattern, and the boundary position was determined using a sliding window of three cells and a majority rule to associate each cell to the most prevalent state in its window (the boundaries are then set to the first position where cell states change from blue to white, and from white to red). (c) Examples of simultaneous growth and patterning under the Mixed, Landscaping, and Bubble rule, respectively, for two different growth rates (cell division probability per time step). (d) Fitness as a function of final system size for the three rules and additionally a modified version of the Landscaping rule (see main text). The vertical dashed lines mark the system sizes beyond which the average time of pattern formation (fig. S1) is expected to exceed the time required to reach a growth speed of one cell per time step. (e) Fitness as a function of the probability of asynchrony *p*_*A*_ for the same four rules. (f) Examples of the patterning dynamics at a probability of asynchrony *p*_*A*_ = 0.1 in a system of size *L* = 34.

#### Robustness of patterning with simultaneous growth

Growth via cell divisions is an integral part of many pattern formation processes in developmental biology. We therefore asked to what extent the three patterning strategies of fig. 5 function also in the presence of cell division. To address this question, we introduced a probability *p*_*divide*_ per time step for cells to perform a symmetric division that conserves the internal state (see ‘Model and Methods’). This homogeneous cell division process causes the system to grow exponentially in size over time, during the patterning process (fig. 7c). While exponential growth is not sustainable and real developmental systems limit growth by additional regulation, our simple model suffices to explore how growth generally interferes with the different patterning strategies.

For small cell division rates, all three strategies successfully form French Flag patterns (fig. 7c, left), albeit with stripe widths that are more variable than without growth (fig. 5). This increased variability is to be expected, given the positive feedback that growth exerts on stripe width (a wide stripe extends with a larger instantaneous speed than a narrow stripe). When the cell division rate is increased, patterning following the Landscaping rule is most affected by the interference with growth (fig. 7c, right). In part, this is due to the creation of new bulldozer states induced by the symmetric cell divisions (fig. S4a,b and fig. S3a,b). The rule can be mutated to eliminate this bulldozer creation process, without affecting the underlying patterning strategy (fig. S1, fig. S3c,d).

A second interfering effect of growth on patterning sets in roughly when the instantaneous growth speed exceeds the speed of the patterning process before the final pattern is reached. Since the different rules have different patterning speeds (see above), this occurs first for the Landscaping rule, then for the Mixed rule, and ultimately also for the Bubble rule, as seen in a rapid drop of the average fitness as a function of the cell division rate (fig. 7d). This appears to be a universal limit for any patterning strategy based purely on local signaling, without additional feedback from patterning onto growth.

#### Robustness against asynchronous updates

While there is ample evidence from different organisms that neighboring cells can maintain synchronously varying gene expression levels (***Liao and Oates, 2017***), synchronization will never be perfect, such that the robustness of patterning strategies against asynchronous updates is a pertinent question. To be able to explore the behavior of the three patterning strategies of fig. 5 for different degrees of synchronicity, we introduced the probability *p*_*A*_ for each cell update to be asynchronous (see ‘Model and Methods’), such that *p*_*A*_ = 0 corresponds to the perfectly synchronous updates considered so far, and *p*_*A*_ = 1 to entirely asynchronous updates.

Although the final tripartite pattern is stable also under asynchronous updates, the process of forming the pattern can be disturbed: For all three patterning strategies, the average fitness decreases with increasing *p*_*A*_ (fig. 7e), since the asynchronous updates can significantly distort the relative width of the formed stripes (fig. 7f). On average, the Landscaping rule is most strongly affected, while the Bubble and Mixed rules perform similarly (fig. 7e). The same mutation of the Landscaping rule that improved its robustness of patterning with simultaneous growth, also improves its robustness against asynchronous updates (fig. 7e). However, even with this mutation the observed general trade-off remains: The Landscaping strategy displays the highest patterning accuracy under perfect conditions, but is sensitive to perturbation by asynchronous updates.

### Tunability of the patterning process

Another relevant set of questions about the patterning strategies of fig. 5 concerns their generality: Can the same underlying strategies also be used to form other axial patterns? How can the quantitative characteristics of a pattern be tuned? And to what extent is the patterning process controlled by the initial conditions or by the update rule, respectively?

As initial conditions we assumed random initial states so far, but another natural choice, inspired by developmental biology, is to assume an “undifferentiated” initial cell state (black) for all cells within the system (i.e., not on the boundary). During the pattern formation process, their cell fate is then ultimately determined to be one of the differentiated states, i.e., blue, white, or red for the French flag pattern. Since the Landscaping rule of fig. 5 implements an erase-and-reconstruct patterning strategy, it should be able to form the French Flag pattern starting from any initial condition, and thus also from the homogeneous undifferentiated pattern. This is indeed the case (fig. 8a). However, other strategies are also available, as revealed by an evolutionary search (fig. 8b,c). For this evolutionary search in 4-state rule space, we adapted the method of fig. 2a by inserting only random rules that satisfy the steady-state conditions of fig. 3c. Note also that the evolutionary search with a homogeneous initial pattern is significantly faster than with random initial patterns, since the average over different initial conditions is not required for the fitness calculation.

**Figure 8.**
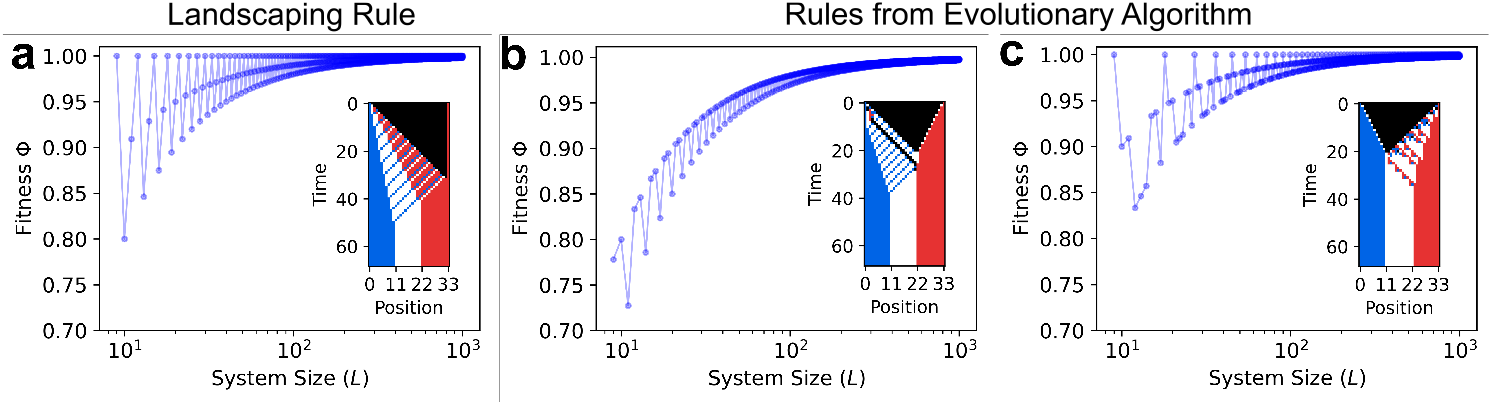
Undifferentiated initial condition. Here, the black state is used as an undifferentiated state, in which all cells within the system are initialized (except for the fixed boundaries). (a) The Landscaping rule solves the French flag problem also for this initial condition. (b, c) An evolutionary search yields a different patterning strategy of two fronts starting from the boundaries and moving inward at speeds of 1 and 1/2 cells per time step, respectively, with the faster front emerging either form the left (b) or the right (c).

Both shown examples (fig. 8b,c) use two patterning fronts that propagate from the boundaries inward. The two fronts move at different speeds, with a 2:1 ratio, and arrest when they collide. The collision point thus either lies at one third of the system size (fig. 8c) or at two thirds (fig. 8b). The slower front generates one stripe, while the faster front generates two stripes in its wake, such that the tripartite pattern ultimately emerges with approximately equal stripe widths. Stripe formation behind the faster front involves a sorting process in both examples, but the detailed characteristics of this process are visibly different (fig. 8b,c).

How can the relative widths of the stripes in the French flag pattern be tuned? To explore this question, we first performed an evolutionary search in 3-state rule space for rules that form the pattern with a stripe width ratio of 1:2:1 starting from uniformly distributed random initial conditions. This revealed a consensus rule that achieves the targeted ratio well for larger systems (fig. 9a,b). This rule uses the relevant entries of a sorting module for red and white cells, while the bulldozer module is changed. The primary effect of this change is revealed by initializing the pattern formation process with isolated blue states in an all-white region (fig. 9c): single blue cells travel from right to left, producing white and red cells in a 2:1 ratio in their wake. We also performed an evolutionary search for 3-state rules achieving other target ratios, 2:1:1 and 3:1:1, for the stripe width fig. S1, which suggests that simple integer stripe width ratios are attainable with similar mechanisms. However, it seems clear that the stripe widths cannot be continuously tuned in this way.

**Figure 9.**
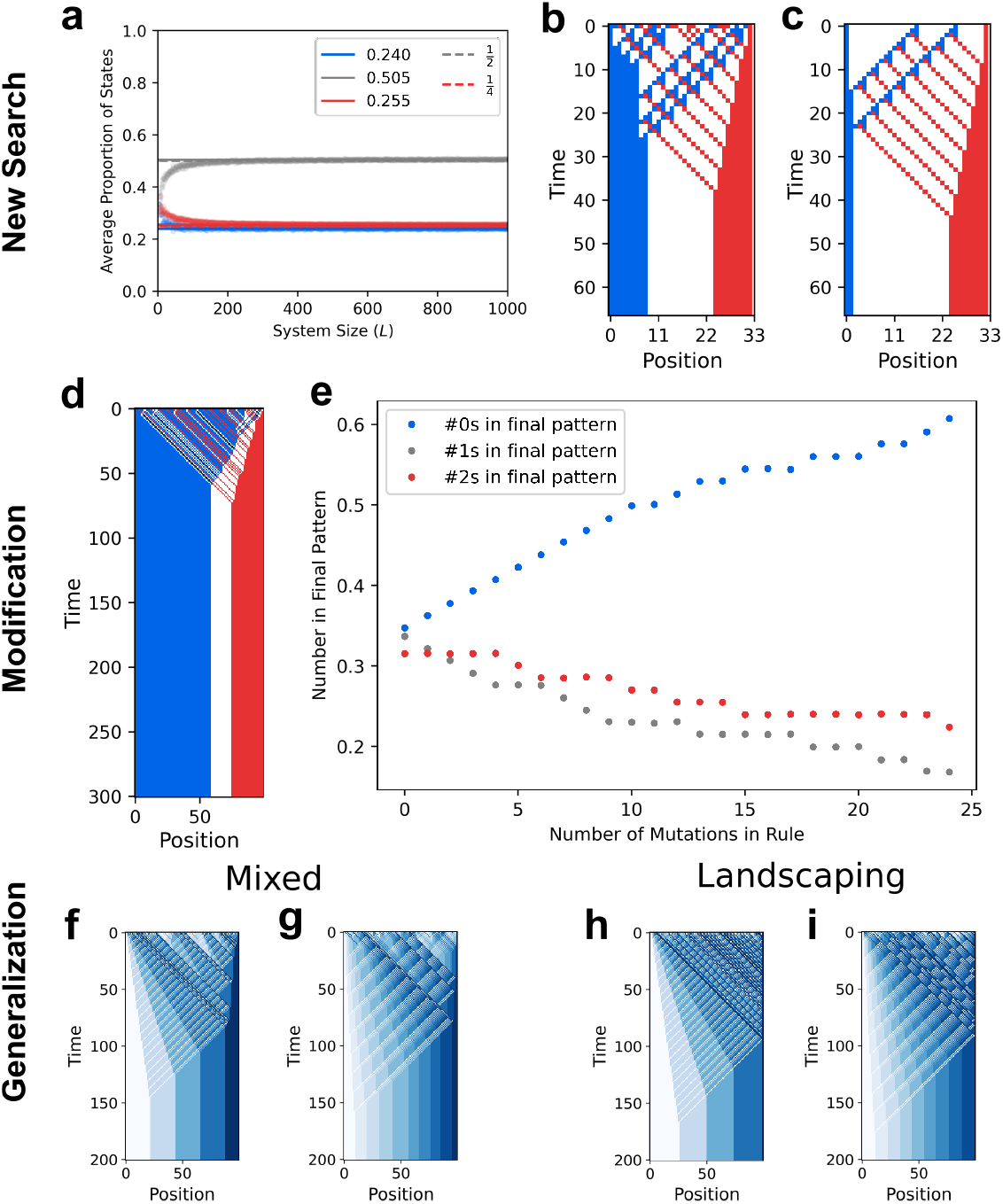
Tunability of the patterning process. (a-c) An evolutionary search for rules that form a French Flag pattern with a stripe width ratio of 1:2:1 from random initial conditions identified a rule (genotype 201 220 122 202 110 112 201 110 000), which (a) achieves this target ratio increasingly well for larger system sizes (dots show the average proportion of each state, dashed lines the aimed ratios, and solid lines the average value from *L* = 901 to *L* = 1000). (b) The patterning behavior of this rule is similar to that of the consensus rule, however (c) the bulldozer state (blue) produces white and red cells in a 2:1 ratio in its wake, as it travels from right to left (shown here using an initial condition with only two isolated blue cells in a white background). (d-e) Mutations of the Bubble rule can tune the resulting stripe widths in fine-grained steps, for instance to (d) increase the width of the blue stripe, in (e) 24 mutations that gradually increase the production of blue states from black states. (f-i) Generalization of the mixed and landscaping strategies to form axial patterns with more than three stripes, as illustrated here using either 5 or 10 states.

For continuous tuning of the relative stripe widths, we turn to the 4-state Bubble rule of fig. 5. A possible way to control the relative width of the stripes for a pure sorting strategy is to control the statistics of cell states in the initial pattern. Indeed, for initial patterns without black states, the Bubble rule conserves the number of blue, white and red states (fig. S2), such that any intracellular mechanism creating a bias in the initial cell state choice will permit the formation of arbitrary relative stripe widths. If black states are included in the initial condition, they are converted to blue, white, or red states by the Bubble rule. Since the resolution module fixes only a fraction of rule entries out of all entries involving a black state (fig. 4), the remaining entries can be modified to bias the creation of certain cell states. This allows to tune the resulting stripe widths in fine-grained steps, for instance to increase the width of the blue stripe (fig. 9d,e), or to increase the width of the white or red stripes (fig. S3).

Finally, we considered target patterns with a larger number of stripes. While the patterning strategy underlying the Bubble rule could, in principle, be generalized to more states, and thus more stripes, this would require a number of additional transient states for resolution modules of the type shown in fig. 4. The generalization of the patterning strategies underlying the Mixed rule and the Landscaping rule is more straightforward, without a growing “overhead” of additional transient states. Utilizing the state of the last stripe (Mixed) or an additional state (Landscaping) as Bulldozer module and sorting the seeded states with pairwise Sorting modules, generalizations of the Mixed and the Landscaping rule are indeed able to form axial patterns with larger numbers of stripes (fig. 9f-i).

## Discussion

We revisited Wolpert’s French flag problem (***Wolpert, 1968***) to identify alternative axial patterning strategies, which do not require long-range signaling, but are strictly based on local, nearest-neighbor interactions. Local patterning strategies can function even when the transport of signaling molecules, for instance via diffusion, is restricted or too slow for effective long-range signaling (***Dickmann et al., 2022***). To describe such strategies, we used cellular automata as a minimal modeling framework that allows to enumerate all possible update rules for the patterning dynamics, and thus all patterning strategies, for a given number of internal cell states (fig. 1). As in the original French flag problem, the modeling framework considers cells that do not move, but maintain their relative arrangement. However, when we searched the model space with an evolutionary algorithm (fig. 2a), one component of successful patterning strategies turned out to be a functional module that effectively implements sorting of cell states (fig. 3d). Rather than physical sorting of cells via differential adhesion (***Steinberg, 2007***), this module sorts the information contained in the cell states, which in biological cells is reflected by their gene expression patterns. This informational sorting is implemented via local cell-cell communication according to the signaling logic specified by the entries of the cellular automaton update rule (fig. 3d).

The pattern formation process can either make use of the initial cell states, or ignore them completely by erasing the initial pattern with a propagating front that generates the intended pattern in its wake (fig. 5). This latter “Landscaping” mode of patterning is thus suitable also when all interior cells of the system are initially in the same undifferentiated state. Patterning can then start from either boundary of the axial system, or from both boundaries simultaneously (fig. 8). However, in a system without global signaling, it is not clear how two patterning waves could be triggered simultaneously from different ends of the system. In contrast, local synchronization of cells, as assumed by the cellular automata modeling framework, is attainable with local interactions (***Uriu et al., 2021***). Additionally, we found that the patterning strategies do not immediately become dysfunctional with a small degree of asynchronicity (fig. 7e). It is also important to note that in order for the patterning strategies discussed here to be viable, cells need to be able to differentiate between left and right neighbors in their signal detection and processing, which requires cell polarization. Although we have limited our analysis to one-dimensional models here, extending the analysis of axial patterning in two- or three-dimensional systems would not pose conceptual challenges. Additional lateral signaling in two- or three-dimensional systems could also be leveraged to increase the robustness of the patterning process to noise, via a positive reinforcement or majority voting scheme (***Ramalho et al., 2021***).

A valuable tool in our analysis of patterning strategies has been the multiple alignment and consensus procedure (fig. 2b), which is enabled by the discrete nature of cellular automata models. This procedure leads to sequence logos and the identification of relevant entries within update rules (fig. 3a,b), and helped us to discover the two basic patterning modules that form the core of the patterning strategies analyzed here: A sorting module that generates patterns by segregating two states away from each other, and a bulldozer module that generates a patterning wave (fig. 3d). We then analyzed the trade-offs between the different patterning strategies: erasing and recreating a pattern from scratch is more accurate (fig. 6), but sorting the initial condition is faster (fig. S1) and more robust to noise and growth (fig. 7). Additionally, sorting is more plastic in the sense that stripe widths can be easily tuned by either biasing the initial condition or mutating a few entries of the rule itself (fig. 9). This is analogous to changing the stripe width in Wolpert’s morphogen mechanism by altering the threshold concentrations for cell differentiation.

Systematic searches for mechanisms for the robust and tunable formation of different patterns, potentially in several spatial dimensions, remains an interesting challenge for the future. Our approach of using evolutionary searches and a multiple sequence alignment procedure to identify functional signaling logics may prove fruitful in this direction, although enlarging the potential rule space might entail the need to employ more complex evolutionary search algorithms ensuring a large solution diversity (***Cully and Demiris, 2018***).

## Acknowledgments

We are grateful to members of the Gerland group and Thiago T. Varella for helpful discussions. G.V. would like to thank the Streicker International Fellows Program for the generous support through the Streicker Fellowship. This work was funded by the Deutsche Forschungsgemeinschaft (DFG, German Research Foundation) – Project-ID 201269156 – SFB 1032 and under Germany’
ss Excellence Strategy – EXC 2094 – 390783311 through U.G.

## Appendix 1 Tables and figures

### Figure 1 - Supplement algorithm S1

Cellular Automata Simulation

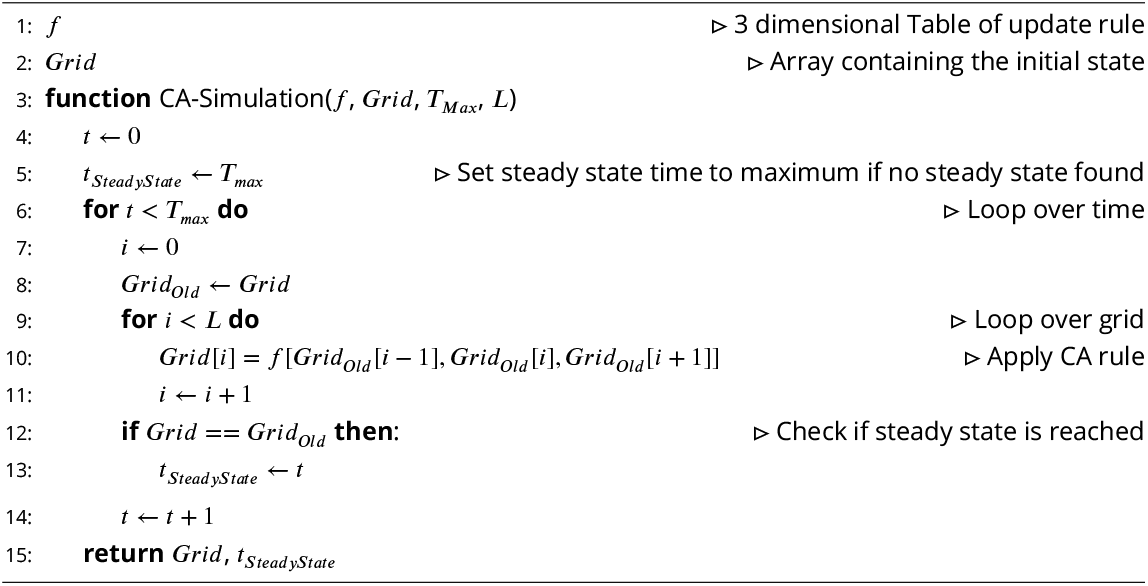

### Figure 1 - Supplement algorithm S2

Fitness Calculation

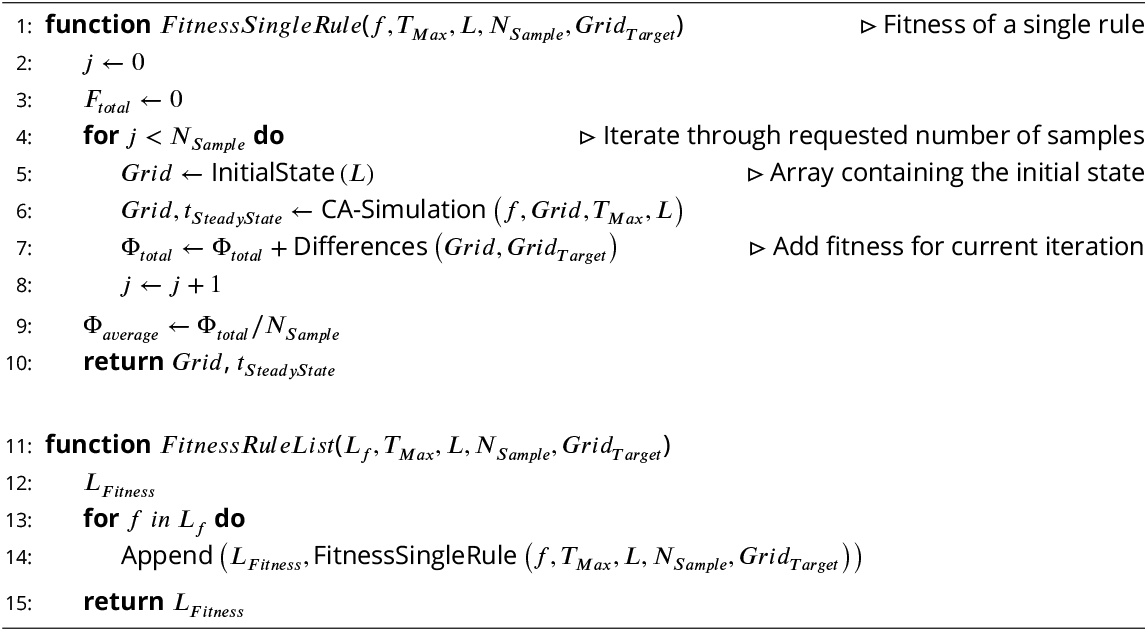

**Figure 2 - supplement figure S1.**
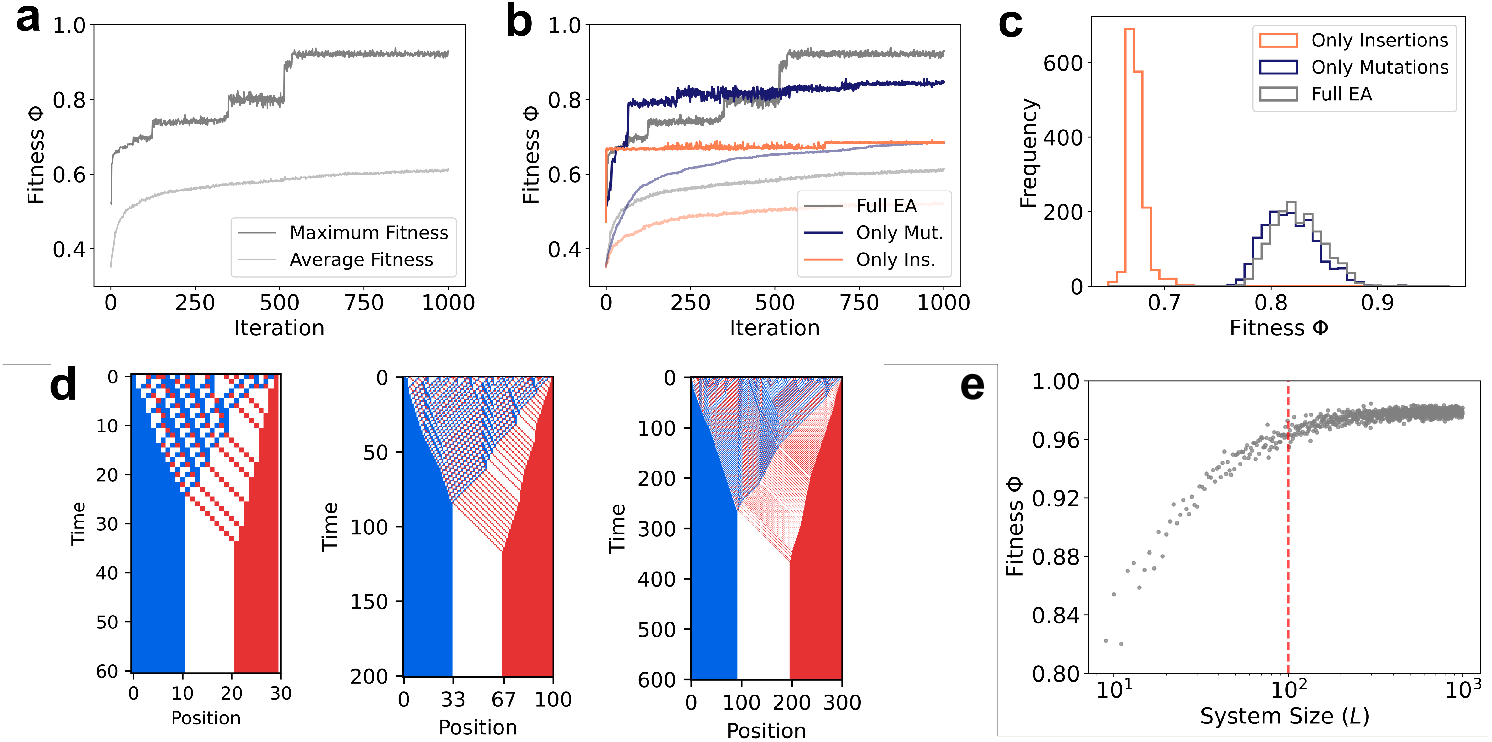
Results and analysis of evolutionary algorithm. For all simulations in this figure, the algorithm was started with 100 random rules and ran for 1000 iterations. Rule fitness was estimated by an average over 50 random initial conditions with CA size of *L* = 100. Each iteration, rules were mutated and, if less fit than the mutant, replaced by the mutant rule. Then, the 30 least fit rules were substituted by random ones. (cf. alg. S1) (a) Average and best fitness of the 100 rules in the list at each iteration as a function of evolutionary algorithm iterations. (b) Comparison between the ‘full’ EA using both mutations and replacement by random rules to modify the rule list, as well as modifications of the EA either only mutating the rules or only randomizing the least fit ones. (c) Distribution of maximum rule fitness found by the algorithm over 1619 runs (1620 for only mutation of rules, 1621 for only insertion of random rules) with the parameters specified before. (d) Phenotype of the fittest rule at the end of the run in (a), also fittest rule of the Full EA distribution in (c), for length *L* = 30, 100, and 300. (e) Dependence of the fitness of the rule in (a, b, d) on system length *L* for an average over 50 random initial conditions as in (a, b, c). Length *L* = 100 for which the fitness is usually calculated is marked with the red, dashed line.

**Figure 2 - supplement figure S2.**
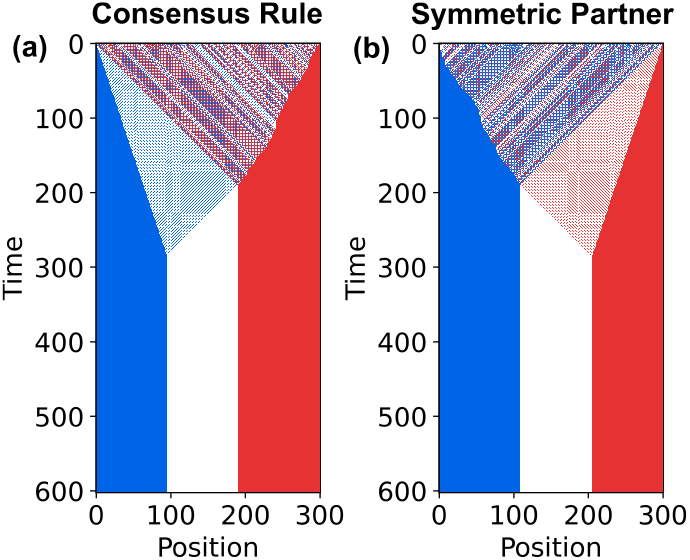
Kymographs of consensus rule (a) and its symmetric partner (b) with genotype *S* = 000011012121011012111022002.

### Figure 2 - Supplement algorithm S1

Evolutionary algorithm simulation

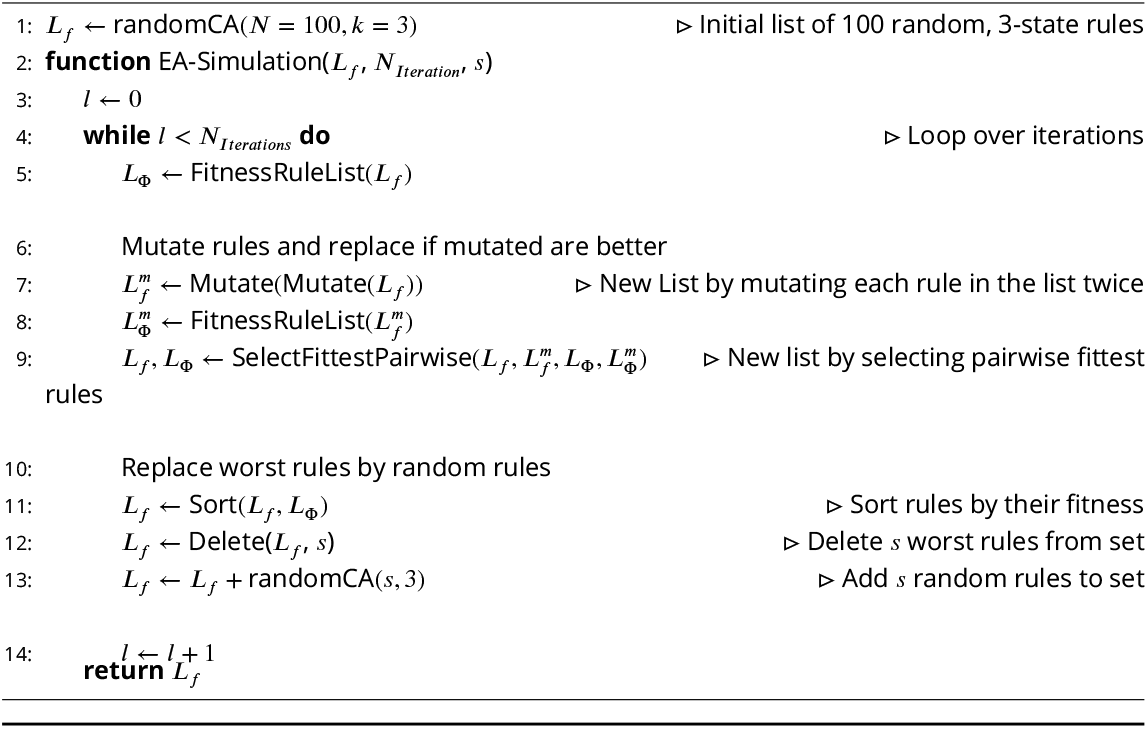

**Figure 2 - supplement table S1.**
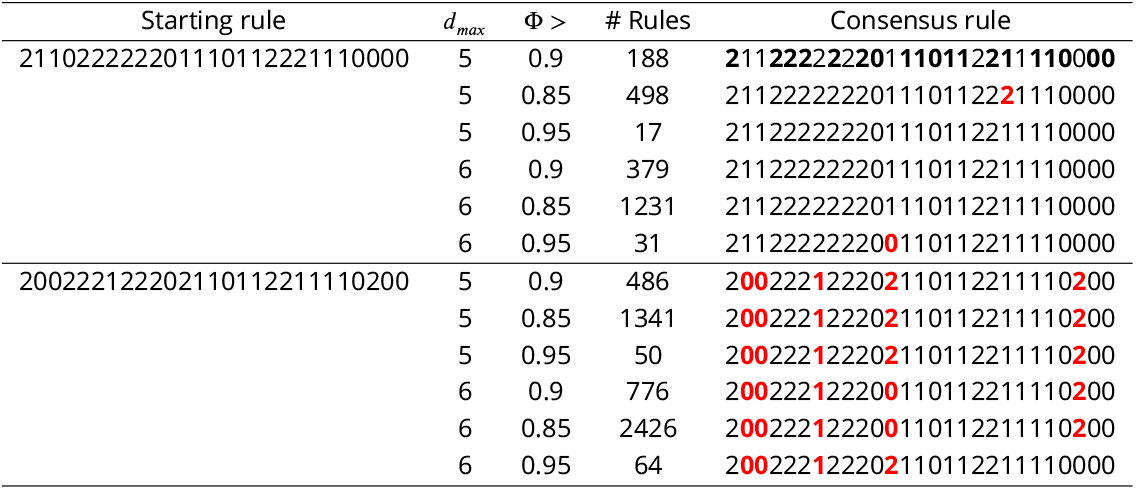
Consensus rule for different parameters. The consensus rule was calculated for 2 different 2 different neighborhood sizes and 3 different fitness thresholds. The differences to the consensus rule in the marked in red, with all consensus rules differed from the one selected by at most 5 mutations.

**Figure 3 - supplement figure S2.**
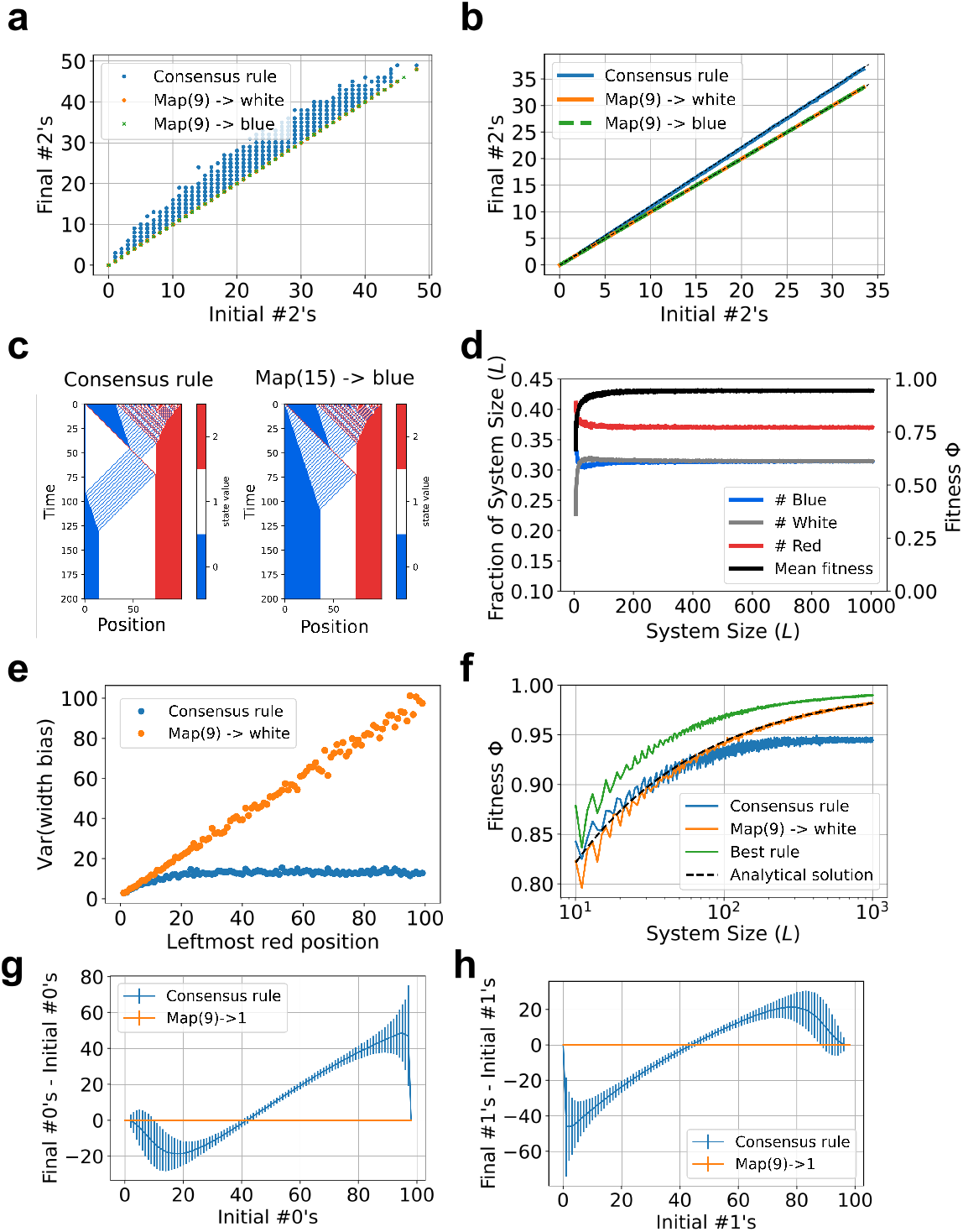
Properties of consensus rule and its modifications. (a) Number of red states in final pattern as a function of the number of reds in the initial pattern for consensus rule, consensus with mapping 9 changed to white and to blue. The number of final reds for the consensus rule is always slightly larger or equal than the number of initial reds, while for both modifications of the consensus the numbers are the same. (b) Same data as in (a) but averaged for each system length. The dashed black lines in the figure are the lines *y* = *x* and 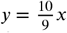. (c) Example of how the bulldozer state in the consensus rule sometimes seeds more whites than blues and how modifying the output of mapping 15 to blue makes the seeding ratio 1 *z* 1. (d) Scaling of the fraction of blue, white, and red states with system length. Mean fitness is also plotted in black. (e) Dependence of the variance of the final difference in the number of whites and blues on the position of the leftmost red state. Each point is the variance over 1000 random initial conditions. For the consensus rule, this variance is capped because mapping 9 introduces a red state close to the left boundary even if there isn’t one in the initial state. Mutating this mapping makes the variance scale linearly with red state position, consistent with the widths being independent binomial random variables. (f) Scaling of the mean fitness with system size *L* for the consensus rule, its modification, and the fittest rule (rule 200 222 122 202 110 112 211 110 100). While the consensus fitness converges to 17/18, the fitness of the modified and the fittest rules converge to 1 as *L* → ∞. The analytical solution plotted is the equation 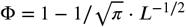. (g,h) Mean difference between the final and initial number of 0s (g) and 1s (h) as a function of the initial number of 0s (g) and 1s (h), respectively. 10000 samples of initial conditions with only blue and white states with a proportion given by the value on the horizontal axis and random sequence. Error bar corresponds to the standard deviation of the sample of each point

**Figure 3 - supplement figure S3.**
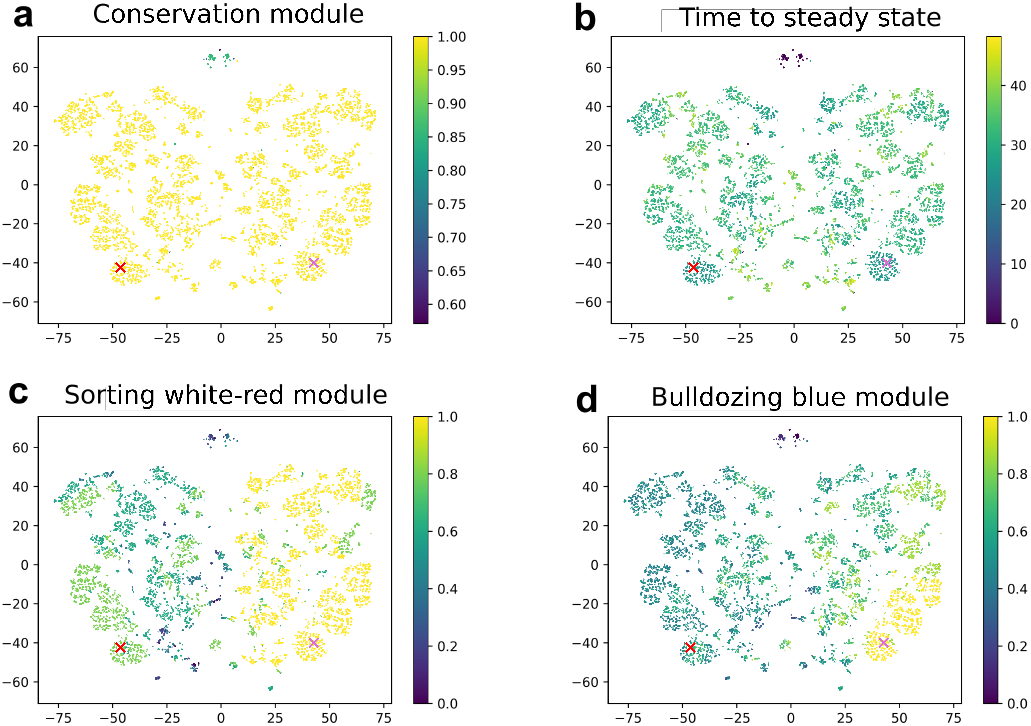
tSNE Plot of all rules with fitness larger 0.85 as described in fig. 3 and Fitness. (a) Number of rule entries that are the same as (b) Time to steady state, (c) Relevant entries (entries with information content larger *log*_2_(3) − 1 bits (cp. fig. 3 b) for a module that sorts red and white cells or (d) a blue bulldozer state moving to the left

**Figure 4 - supplement figure S1.**
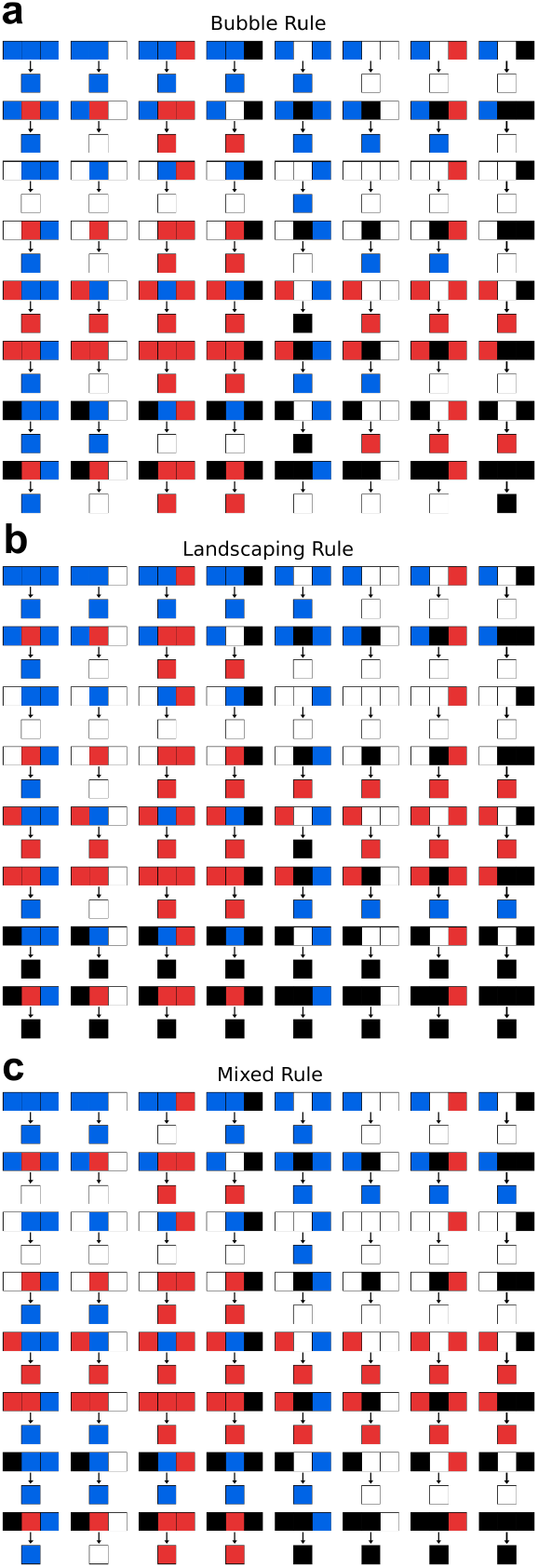
Complete rule tables for the engineered rules in the *k* = 4 space.

**Figure 4 - supplement figure S2.**
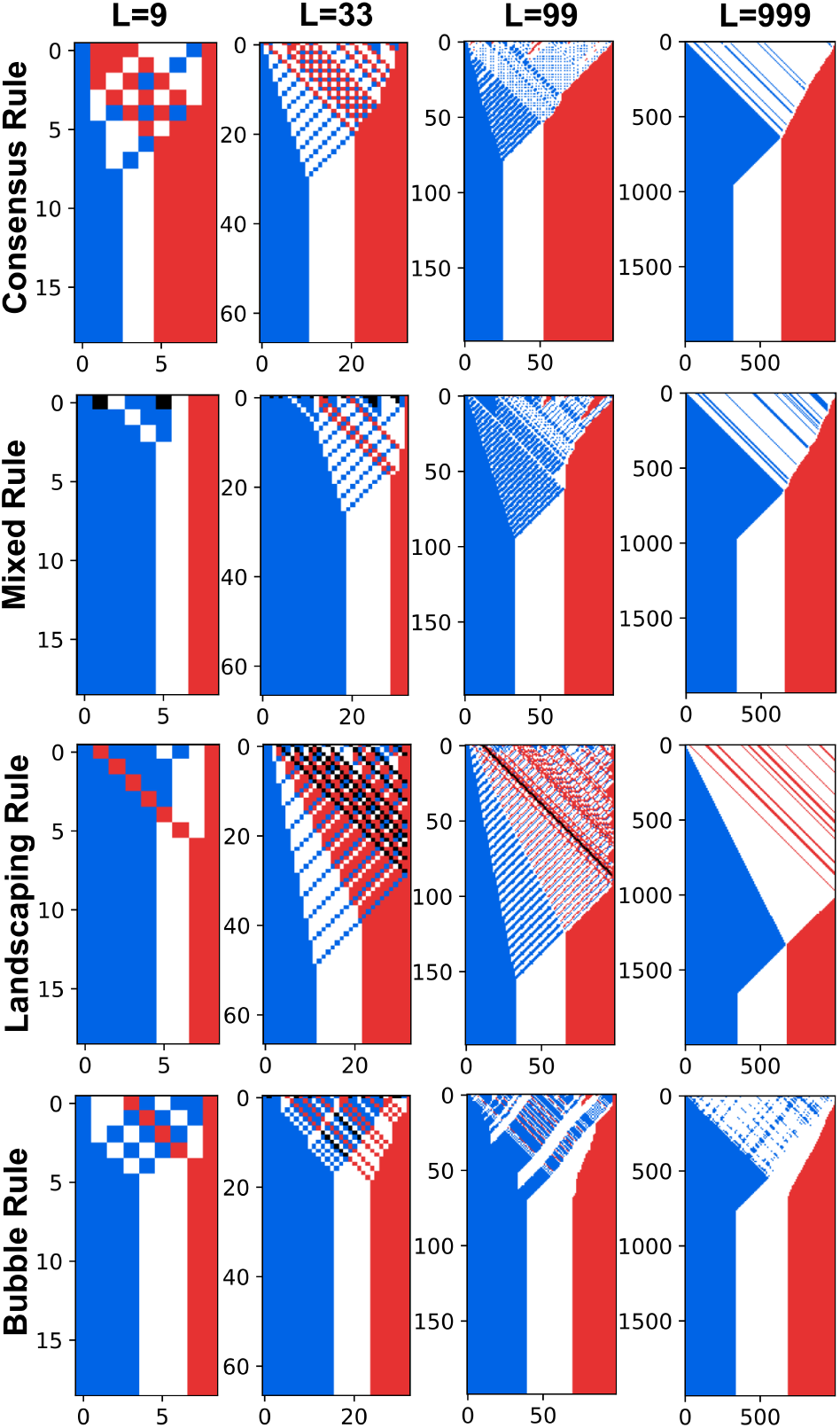
Simulation examples of the prototypical rules for different system sizes *L* ∈ {9, 33, 99, 999}

**Figure 6 - supplement figure S1.**
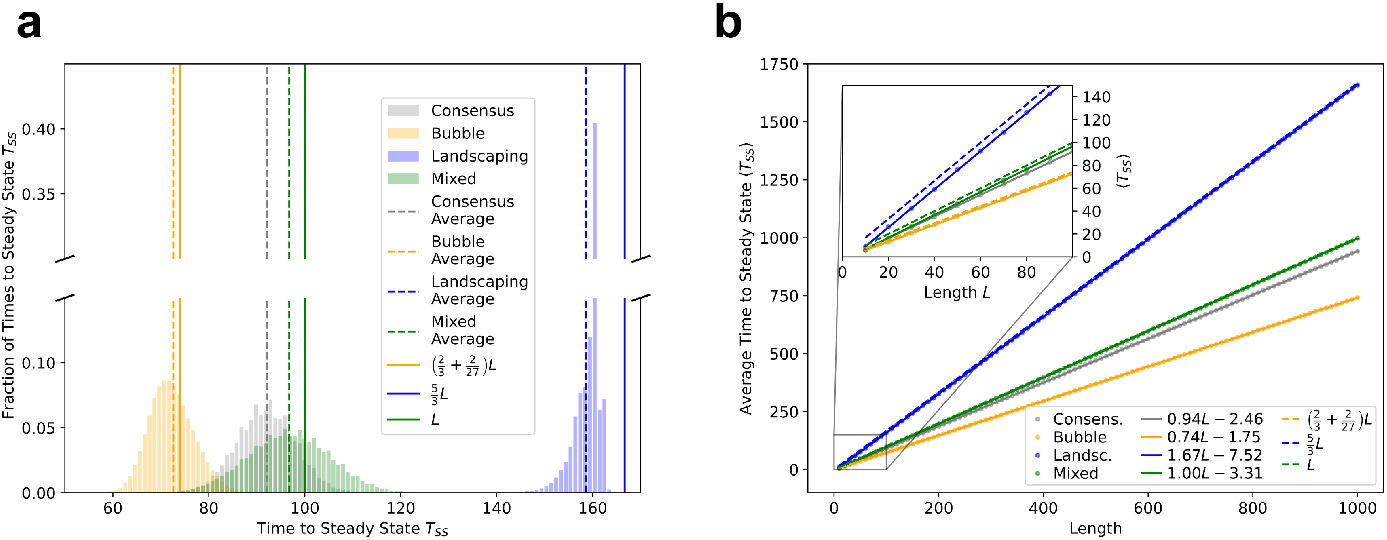
Steady State Times for Consensus, Bubble, Landscaping, and Mixed rule. (a) Sampled distributions for *L* = 100 with the rough estimates of the average from the main text. (b) Scaling of average steady state time with length (dots), together with the rough estimates (dashed lines) and the fits (solid lines).

## Appendix 2 Choice of fitness function

The algorithm to calculate the fitness for a single rule according to eq. (3) is outlined in Fitness Calculation. (For fig. 7a, *N*_*sample*_ = 100, *T*_*max*_ = 2*L*). The fitness serves two separate purposes in our argument: as a measure of how well a particular rule is able to generate a specific target pattern, and as a score function in the evolutionary algorithm.

Depending on the target pattern and the (biological) context, different measures might be used to better capture the first purpose, e.g., for the French flag pattern a cell with a red state where a blue cell should be situated could be punished more severely. Here, the Hamming distance was chosen as fitness because it is independent of the specific target pattern and arguably the simplest possible choice.

In it’s function as a score for the evolutionary algorithm, a different choice of fitness function might be useful to speed up the convergence of the EA algorithm by, e.g., introducing a non-linearity before summing up the individual scores for different initial conditions. We did not use these possibilities here to keep a single consistent fitness function throughout the whole paper. Our choice of fitness function is, for well working cellular automata rules, usually not constant with system size but a mostly increasing function of the system size *L* (fig. S1 d, e), depending on the sample size and the exact value of *L*, since only every third size is divisible by three as needed to for an unambiguous choice of target pattern for a French Flag pattern.

When searching rules that form other target ratios also different length (*L* ∈ {40, 100, 200}) and simulation times (either *T*_*max*_ = 2*L* or with equal weights *T*_*max*_ = 2*L* and *T*_*max*_ = 5*L*) were used to counteract the fact that found solutions only worked for specific lengths and/or a specific simulation times.

## Appendix 3 Evolutionary Algorithm

The basic evolutionary algorithm as conceptually shown in fig. 2 is represented in pseudo-code in Evolutionary algorithm simulation. The EA is initialized with a list of random rules. A single iteration, which will be carried out *N*_*Iteration*_ = 1000 times, consists of a calculation of the fitness of each rule, as well as mutating each rule twice and calculating the fitness of the mutated rules as well. Each rule is then compared in fitness to its mutated counterpart and the better performing one is chosen. After sorting these rules, the worst 30% are discarded and replaced by random rules before a new iteration starts.

**Figure 7 - supplement figure S1.**
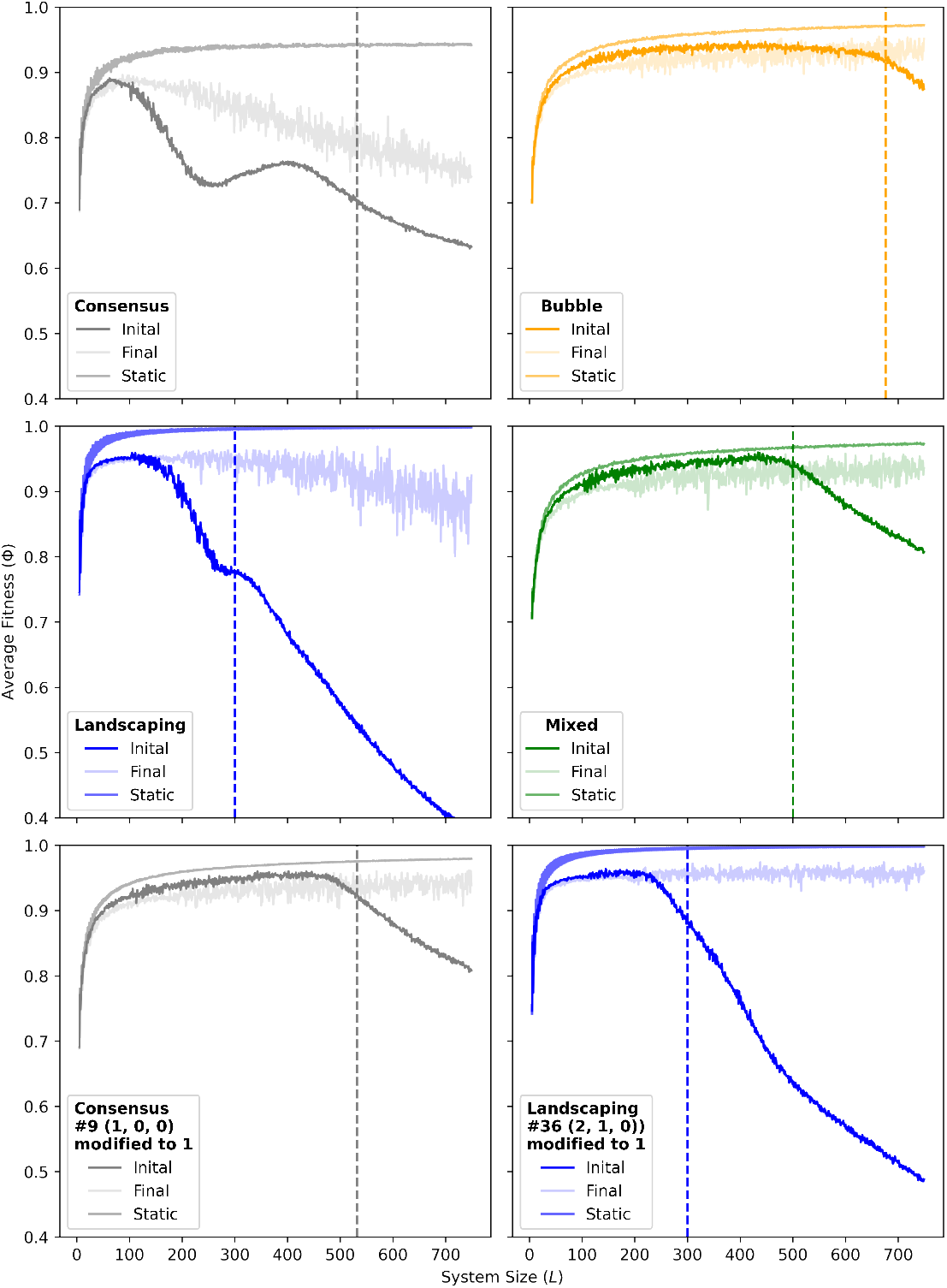
Average fitness of prototypical rules in growing systems. Average fitness as a function of the system size for the prototypical rules in 3 and 4 state space for the Consensus rule, Bubble rule, Landscaping rule and Mixed rule, as well as a modification of the consensus rule in entry 9 to 1 (from 2) and modification of the Landscaping rule in entry 36 to 1 (from 3). In each case, the fitness as a function of the initial and final length are shown for a system growing with a probability of 0.2% per time step per cell. Sample sizes are 10000 for *L* ≤ 10, 1000 for 10 < *L* ≤ 100 and 100 for *L* > 100. Further, the fitness for the corresponding non growing (static) system is plotted. Sample sizes in this case are 10000 for *L* ≤ 10, 1000 for *L* > 10. The vertical lines correspond to the length at which on average the exponential growth of the system exceeds the length at which the exponential growth of the system is faster then the pattern formation process defined by 0.002*aL*_*crit*_ = 1 with *a* being the average number of steps needed to complete pattern formation (cp. S1).

**Figure 7 - supplement figure S2.**
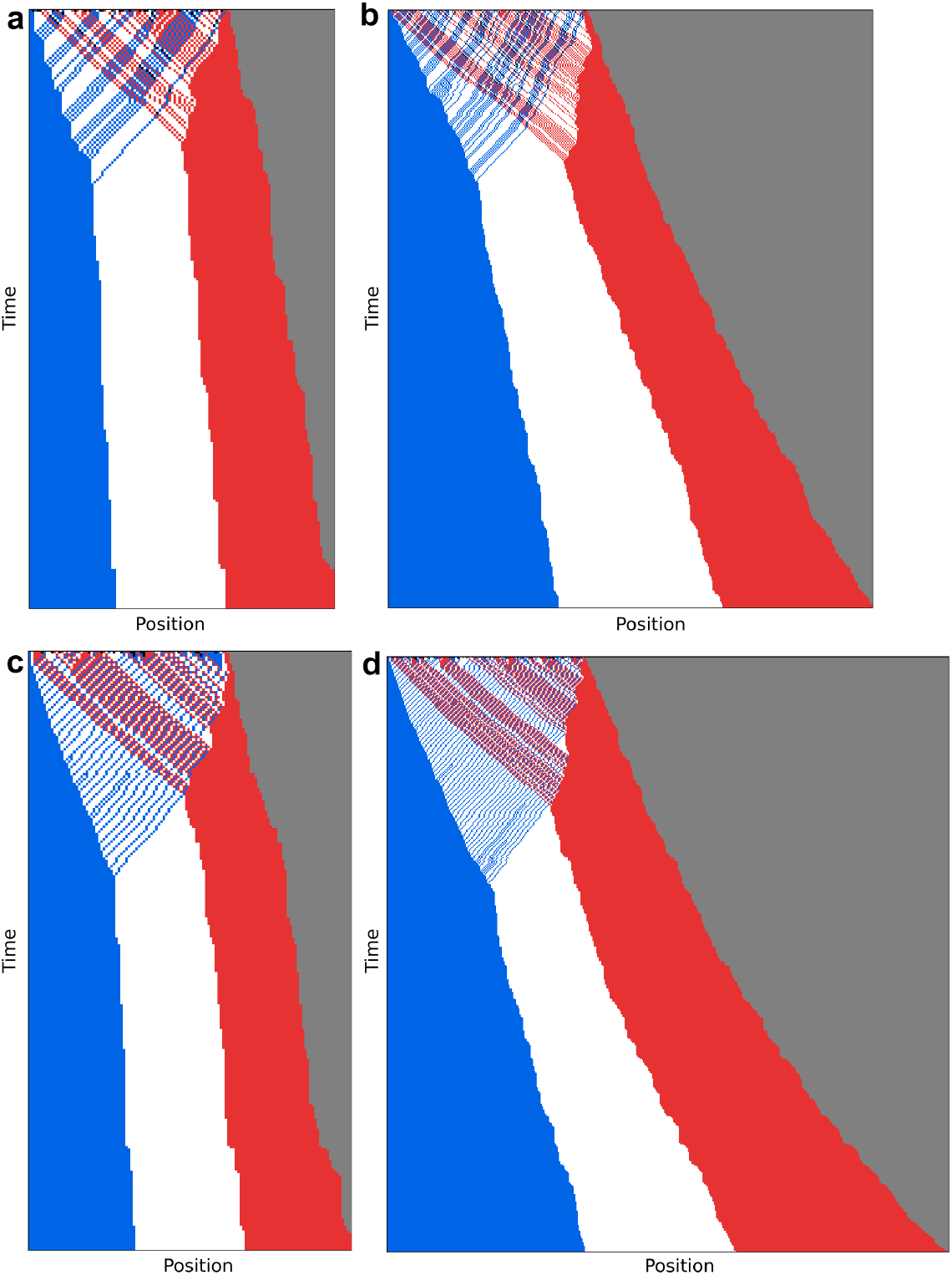
Simulation of a growing system for the Bubble (a, b) and Mixed (c, d) rule for initial system sizes *L* = 80 (a, c) and *L* = 160 (b, d). The system is growing with a division probability of 0.2% per time step per cell.

**Figure 7 - supplement figure S3.**
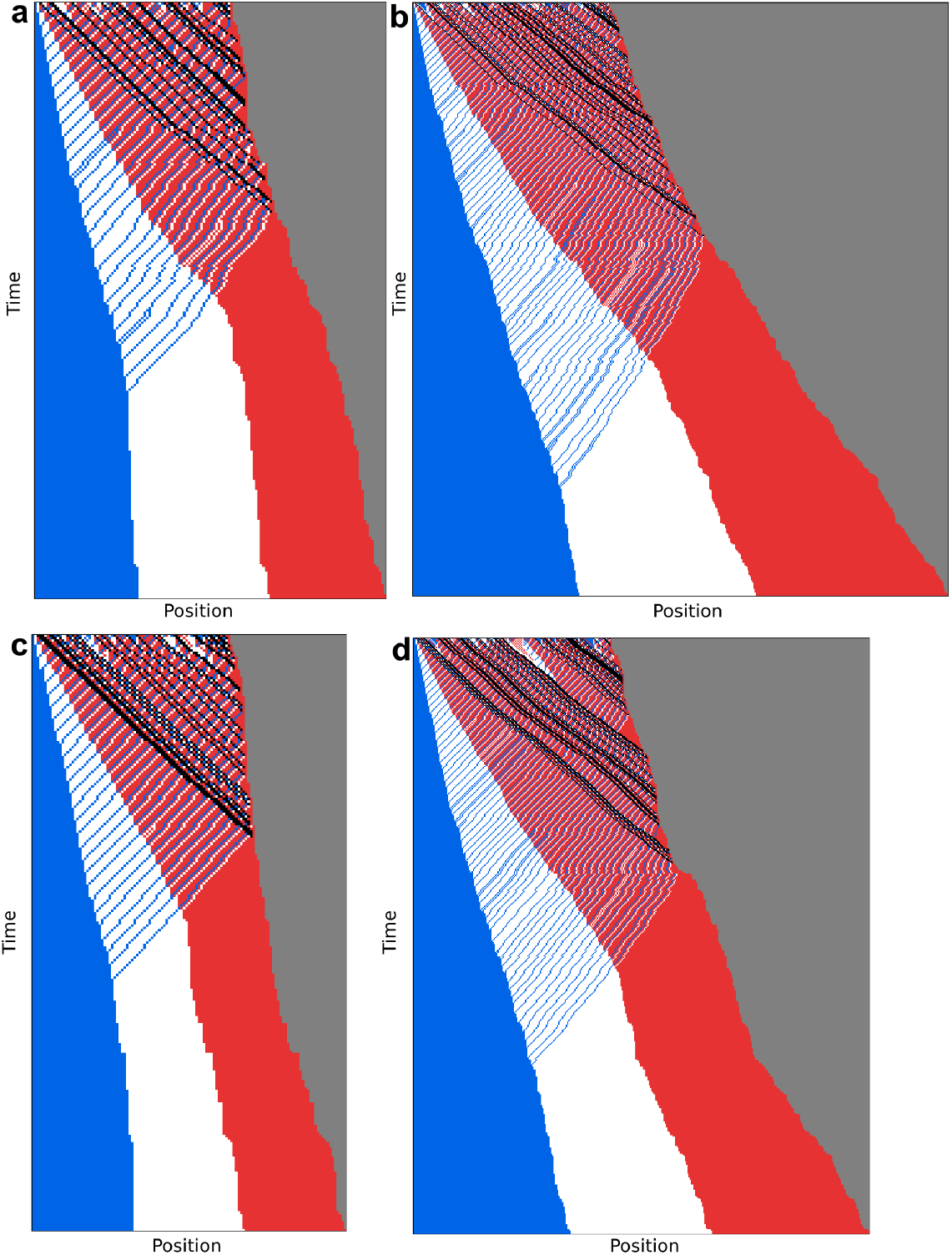
Simulation of a growing system for the Landscaping rule (a, b) and the Landscaping rule modified at position 36 (to 1) (c, d) for initial system sizes *L* = 80 (a, c) and *L* = 160 (b, d). The system is growing with a division probability of 0.2% per time step per cell. The modified entry suppresses the formation of new bulldozer states due to cell divisions

**Figure 7 - supplement figure S4.**
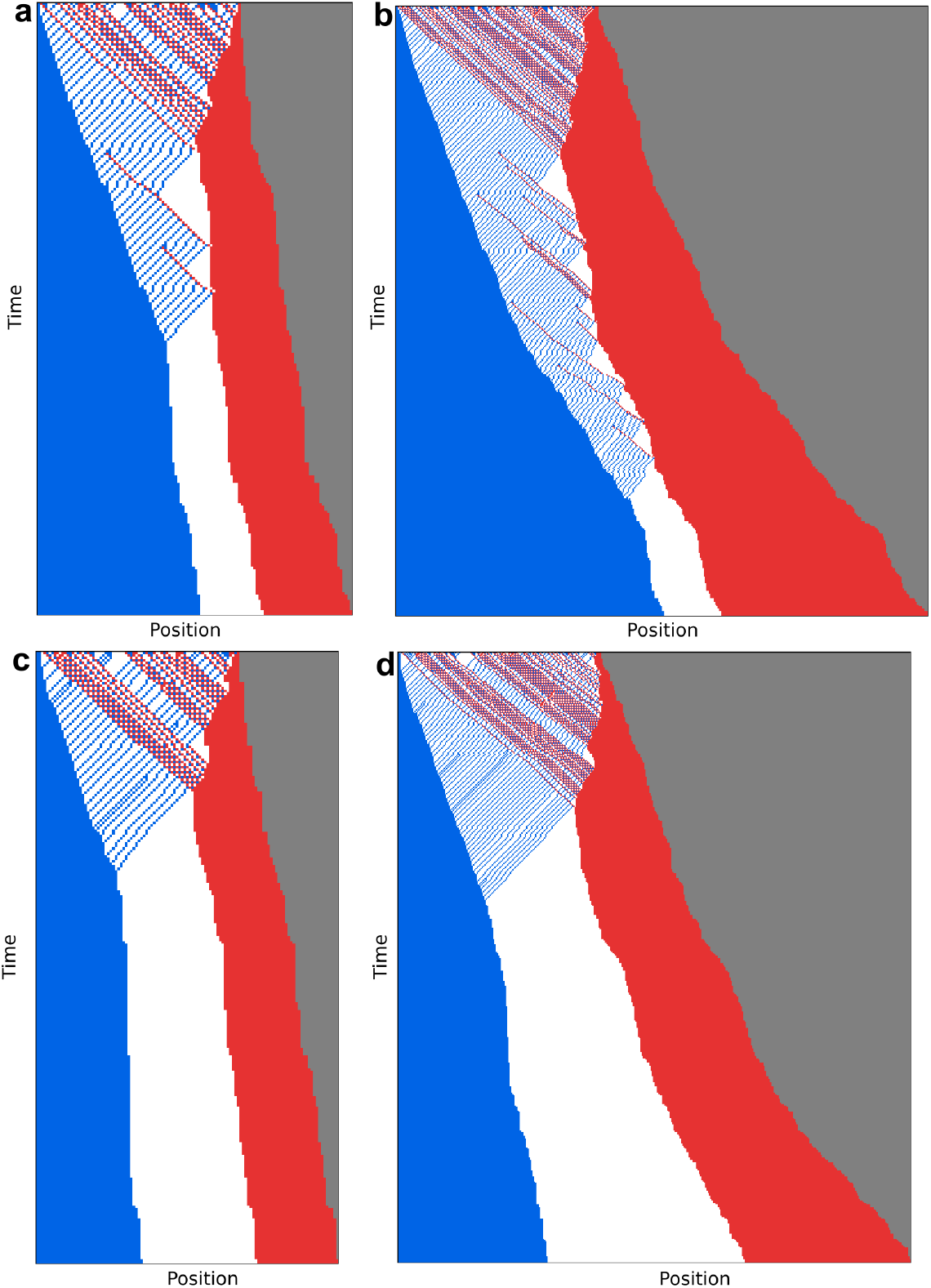
Simulation of a growing system for the consensus rule (a, b) and the consensus rule modified at position 9 (to 1) (c, d) for initial system sizes *L* = 80 (a, c) and *L* = 160 (b, d). The system is growing with a division probability of 0.2% per time step per cell. The modified entry suppresses the formation of new bulldozer states due to cell divisions

**Figure 8 - supplement figure S1.**
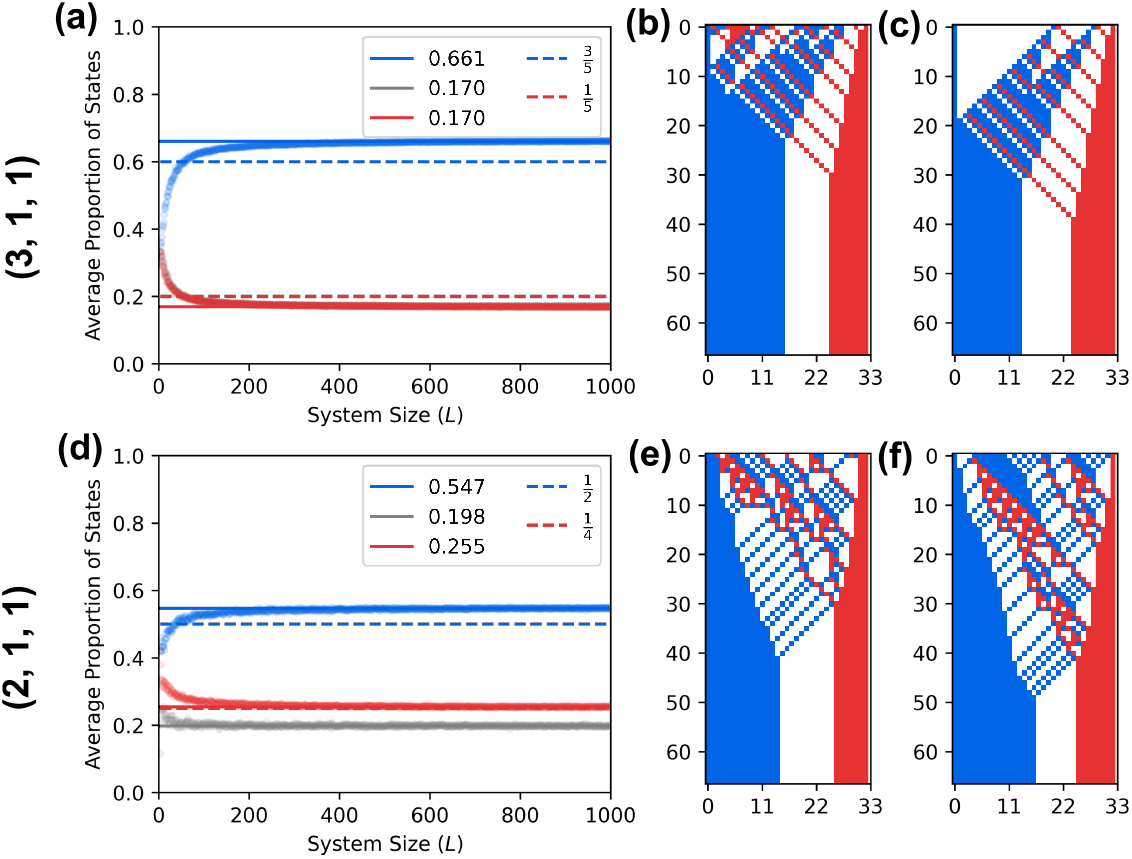
Rules for different target ratios (b, w, r) = (3, 1, 1): 200220122210110102110110100 and (b, w, r) = (2, 1, 1): 211220022210110102111110100. Left column: average proportion of each state (dots) together with the aimed ratios (dashed lines) as well as the asymptotic values (average value from *L* = 901 to *L* = 1000, solid lines). Middle column: exemplary kymograph from random initial conditions for *L* = 33. Right column: from two times adjacent blue and red cells (3, 1, 1), and randomly initial conditions with only blue and white cells (2, 1, 1), respectively.

**Figure 8 - supplement figure S2.**
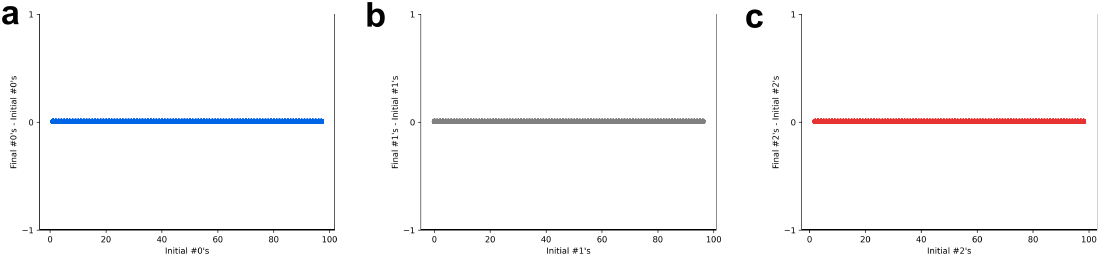
Conservation of States for the Bubble Rule: Difference between number number of states in the blue (a), white (b), and red (c) color in final and initial state when starting from initial conditions without black states for *L* = 99 and 100 randomly drawn samples for every possible total number of blue, white and red initial conditions.

**Figure 8 - supplement figure S3.**
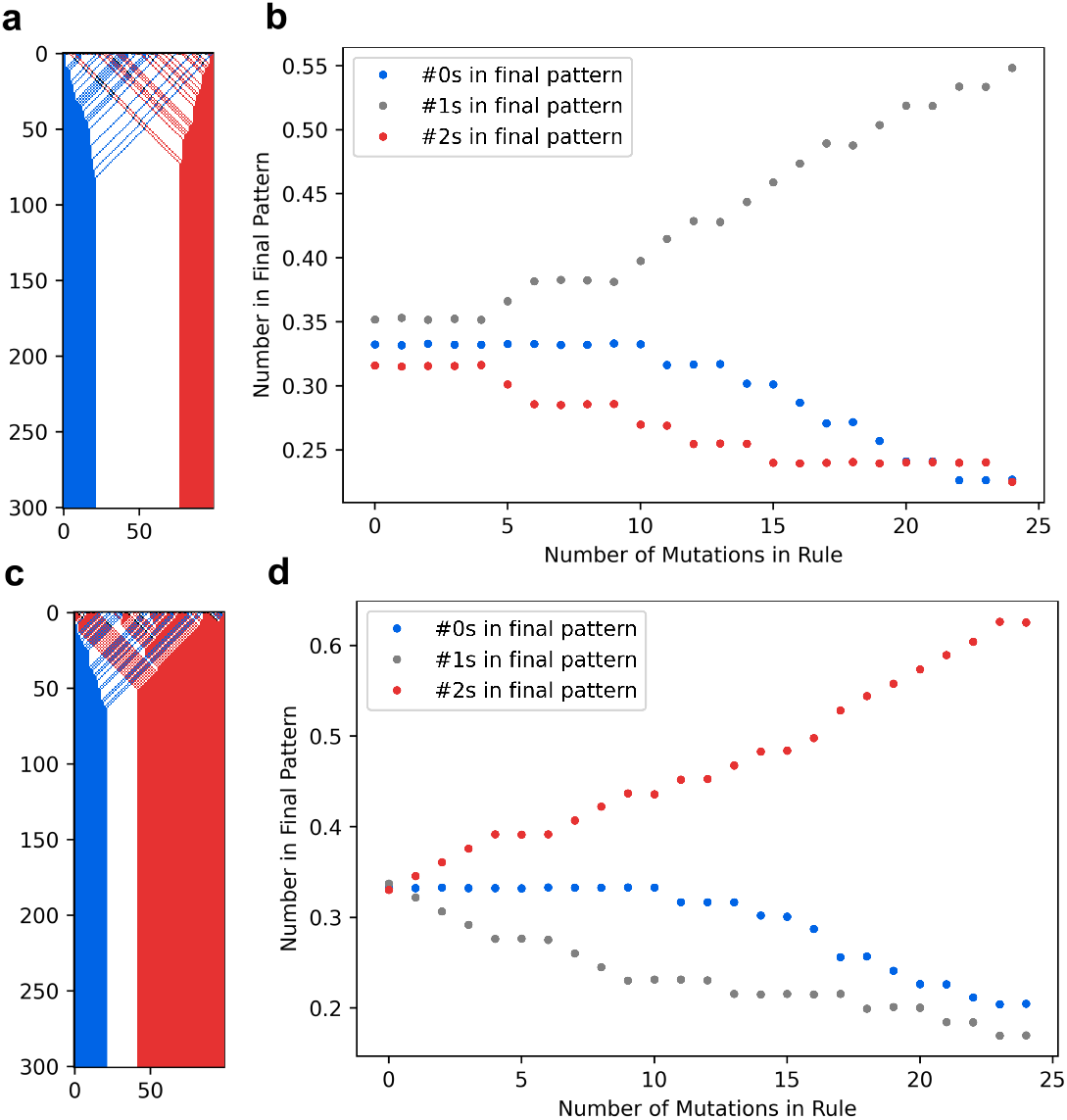
Modifications of the Bubble rule lead to different stripe width. Example (a, c) and average relative stripe width in final pattern as a function of a chosen mutation trajectory (b, d) for modification towards white (a, b) and red (c, d) state, respectively. In each case, the average stripe width can be adjusted using modifications

While most EA runs produce high scoring rules consistently, acquiring exceptionally well scoring rules may need several EA runs (fig. S1 c), which is feasible since a typical EA run was measured to take approximately 4 min on average (on a single core of a Intel Xeon E-2246G 3,60 GHz CPU). While we consider a very specific version of EA algorithms here combining mutations and replacement of the worst rules by random rules, only mutating would also be able to find good rules, although not as quickly as the combination with replacements (fig. S1 b), while just using random replacements (which is essentially equivalent to random sampling) is not able to produce variants with high fitness quickly (fig. S1 b).

## Appendix 4 Computational analysis of the consensus rule

In the main text possible properties of the consensus rule have been hypothesized: (1) the total number of red states is conserved throughout development, (2) the leftmost red state seeds blue and white states at a ratio 1 : 1, (3) the blue and white states to the left of the leftmost red state are sorted into stripes maintaining their numbers, and (4) the pattern formation process scales with the system size. Here, we check these properties computationally for the consensus rule and, when they don’t hold exactly, we explain the mismatch between the consensus patterning strategy and the intuition herein developed and propose modifications to the consensus rule achieving the proposed properties.

The first property, conservation of red states, only holds true approximately for the consensus rule (fig. S2a, b), because the output of its mapping 9 (input: white, blue, blue) slightly increases the number of red states during development. However, mutating this mapping’s output to either blue or white would yield conservation without disrupting pattern formation (fig. S2a, b and fig. 3d). The second property, alternating seeding of blue and white by the leftmost bulldozer state, also only holds true approximately because mapping 15 of the consensus rule (input: white, red, blue) disrupts the perfect alternation between blue and white states (fig. S2c and fig. 3b). Mutating this mapping’s output to blue, instead of white, results in alternating seed without disrupting pattern formation (fig. S2c and fig. 3d).

The third property is also not exact for the consensus rule because mapping 9 prevents blue and white states from just sorting into stripes and can, instead, create red states in between them. Mutating mapping 9 to white would resolve the situation (fig. S2e, g, h).

The fourth property, scaling with system size, holds true for the consensus rule in the sense that the mean position of the boundaries between stripes scales linearly with the size of the system (fig. S2d), a property usually associated with gradient based pattern formation (***Ben-Zvi et al., 2011***).

## Appendix 5 Analytical models

We devised simplified models to describe the patterning strategies of the 3 and 4-state rules. These models have analytically solvable statistical properties and can clarify the connections between patterning strategies and the resulting mean fitness. Moreover, these analyses allow us to quantify to what extent more detailed properties of these rules modify the generic behavior captured by these analytical models.

We model the Consensus rule as a strategy that starts with a random initial condition with white, blue and red states being equally likely. Because mapping number 9 transforms blue states into red states, after the first update the pattern we assume a slight bias towards red, with a probability of 1/3 + 1/27 that an initial state will be red, and with probability 1/3 − 1/27 it will be blue. We make the simplifying assumption that the distribution of these states is still binomial (fig. S1a) and that only the mean of the distribution is changed. After the first rule update, all red states accumulate to the right, moving one position at a time, and maintaining their initial number. In addition, blue and white states to the left of the leftmost red state are sorted and also maintain their number, and finally the states between the initial and final position of the leftmost red state are equally split between blue and white states and sorted.

We model the Mixed rule similarly to the consensus, except that its initial condition includes black states. We assume that these black states get equally redistributed into blue, white and red during the initial updates and then pattern formation continues as in the consensus rule, but with mapping 9 not creating new reds.

We model the Landscaping rule in a similar fashion, but now black states, instead of reds, are used for patterning and the black states disappear from the pattern when they reach the right boundary. We assume all states to the left of the leftmost black state maintain their numbers and are sorted, and the states between the initial position of the leftmost black state and the right boundary are equally split into blue, white and red states.

We model the Bubble rule as a strategy that starts with a random initial condition with white, blue, red and black states being equally likely. As with the Mixed rule, we assume the black states get equally redistributed into the other three states during the first updates. We then assume that the remaining states are sorted, keeping their numbers constant.

Focusing on the fitness function, the models described above always develop into patterns with blue, white and red stripes in this order, but possibly with different widths. To describe these final patterns, we define the variables *b, w*, and *r* to represent the widths of the blue, white, and red stripes in the steady state patterns. The conditions on these widths is that they span the whole flag (*b* + *w* + *r* = *L*) and that they are non negative (*b, w, r* ≥ 0). Using these new variables, we can rewrite the fitness function defined in eq. (3) as:

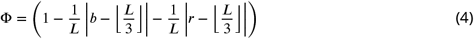

Based on this equation, we calculate how fitness scales with automata length for each of the rule models described above. The variables we need to determine in this calculation are *b* and *r*.

As a first simplification step we assume that *L* is a multiple of 3. The expected fitness value can then be written as:

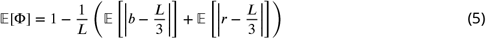

If *L* is not a multiple of 3, the difference to the above expression scales as at most 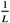

In the Consensus rule model, the number of red states in the final pattern is the same as the number of red states in the initial pattern after the biasing effect of mapping 9. The distribution of *r*, therefore, will be a binomial with parameters *n* = *L* and *p* = 1/3 + 1/27: ***B***(*L*, 1/3 + 1/27). We can use this to derive a lower bound for the last expectation value on the RHS of eq. (5) using the positivity of the absolute value:

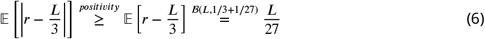

Using the Chernoff-bound for a binomial, we can also arrive at an upper bound for the last expectation value on the RHS of eq. (5):

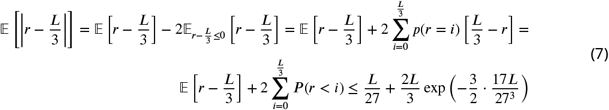

The distribution of *b* is more involved. Assuming the leftmost red state is at position *x* after the effects of mapping 9, the number of blue states between positions 1 and *x* will be distributed following ℬ(*x*, 1/2) and half of the states between positions *x* and *L* − *r* will be blue. Therefore, the distribution of blue states given *x* and *r* can be written as *b* = (*L*−*r*−*x*)/2+*s*, with *s* being distributed as ℬ(*x*, 1/2)

This allows now the calculation of the first expectation in eq. (5). Note that the variables *r* and *x* that are not independent. So, we derive bounds by plugging the above expression for *b* into the expectation and use the triangle inequality as well as the bounds on *x* as a random variable between 0 and *L*:

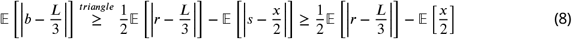

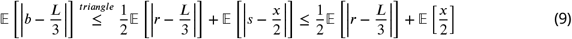

The expectation value of *x* is bounded by the expectation of a geometrically distributed variable with success probability *p* = 1/3 + 1/27 (the probability a state is red) and the other expectation value in the expressions is the one estimated before.

Summarizing everything, we have the following bounds on the difference between 1 and the fitness function:

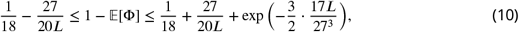

where 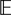[*x*] was approximated by the (slightly larger) expected value of the entire geometric distribution. Therefore, we can conclude that for this model of the Consensus rule,

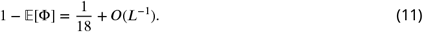

This explains why the fitness of the consensus does not converge to 1: mapping 9’s bias prevents the French flag from developing perfectly even for large system sizes. Mutating this mapping’s output to white instead would solve this problem and we could model this new 3-state rule in the same way we model the Mixed rule next.

In the Mixed rule model, the number of red states in the final pattern is the same as the number of red states in the initial pattern after the redistribution of black states. Since we are assuming black states are redistributed uniformly, the distribution of *r* will be a binomial distribution with parameters *n* = *L* and *p* = 1/3: ℬ(*L*, 1/3) (fig. S1 (b)). Just like before, the distribution of *b* is more involved. Assuming the leftmost red state is at position *x*, the number of blue states *s* between positions 1 and *x* will be distributed following ℬ(*x*, 1/2) and half of the states between positions *x* and *L* − *r* will be blue. Therefore, the distribution of blue states given *x* and *r* can be written as *b* = (*L* − *r* − *x*)/2 + *s*. We again have to calculate the expressions in eq. (5). Since now 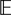[*r*] = *L/*3, the second expectation value on the right hand side is called the mean absolute deviation (MAD) and in this case *r* is a binomial variable, so a closed formula exists. This second expectation does not converge exponentially to its limit value anymore, but rather converges as a power law, as detailed later. We can bound the first expectation in the same way we did before (eq. (8) and eq. (9)), but now knowing that the expectation value of *x* is bounded by the expectation of a geometrically distributed variable with success probability *p* = 1/3 (the probability a state is red). The second expectation value in the expressions is the MAD that already appeared before and can be calculated.

We then have the following bounds on the difference between 1 and the fitness function:

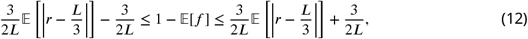

where 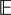[*x*] was approximated by the, slightly larger, expected value of the entire geometric distribution. Using the closed formula for the MAD of a binomial variable (***Diaconis and Zabell, 1991***) and its approximation (***Blyth, 1980***) applied to our specific case, we then have that:

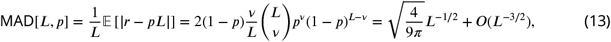

where *v* = ⌊*pL*⌊ + 1. Therefore, we can conclude that for this model of the Mixed rule and for the modified consensus rule,

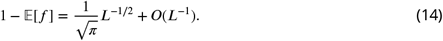

Testing these predictions numerically shows that they work well for the modified consensus but that the constant multiplying *L*^−1/2^ is slightly off for the Mixed rule. We attribute that to the fact that the initial distribution of states is not exactly binomial (fig. S1 (b)) for this rule even though on average the states are still equally distributed forming a French flag and guaranteeing the convergence of this rule’s fitness to 1.

Similarly for the Bubble rule, we now have *b* and *r* distributed as ℬ(*L*, 1/3) (fig. S1 (c)-(e)). Therefore

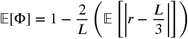

and using eq. (13) and its approximation again, one can show that:

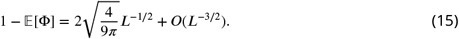

Moving on to the model for the Landscaping rule, we can trace some parallels. Assuming the position of the leftmost black state is *x*, both amounts *r* and *b* have the same distribution and that is *r* = *b* = (*L*−*x*)/3 + *q* with *q* distributed as ℬ(*x*, 1/3). Therefore, plugging this into eq. (4) and taking the expected value we find:

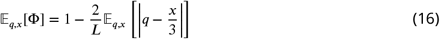

with the indices on the expectation value indicating the variables of the corresponding probability distribution added for clarity. Using a conditional expectation value on *x* we get:

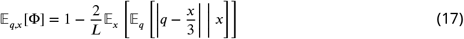

The inner expectation is the MAD of a binomial variable without the normalizing factor of *x*, similar to what we had before. Using the closed formula from before (eq. (13)) for the first term of the expectation, we can get a lower bound to the expression. Additionally, based on the fact that the non-normalized MAD[*x*] is a concave, increasing function in *x*, we can get an upper bound by calculating the MAD for 4, since 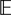[*x*] < 4:

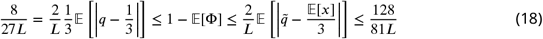

with 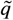 being distributed as ℬ(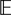[*x*], 1/3). These bounds prove that for this model of the Landscaping rule

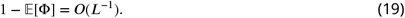

with a coefficient between 8/27 and 128/81.

### Time to steady state

We computationally calculated the scaling coefficients of the mean times to steady state for the 4-state rules described, as shown in fig. S1. Here, we develop an intuitive explanation for their values.

The mean time to steady state of the Mixed rule scales as *T*_*ss*_ ≈ 1.0*L*. This scaling can be understood as the time it takes for the leftmost red state to reach the red boundary (on average 2*L/*3 steps since the state moves one position at a time) plus the time it takes for the last blue state seeded by the bulldozer to sort into place (on average *L/*3 since it is seeded at the red boundary and has to move to the blue boundary).

In the case of the Landscaping rule, the scaling coefficient is 1.67 and a similar intuitive explanation applies. This time, the leftmost black state is the bulldozer and needs to travel the whole length of the system to finish the patterning process. In addition, the last blue state seeded by this hidden bulldozer would need to travel on average 2*L/*3 steps. Overall, then, the pattern formation would take on average 5*L/*3 time steps.

Finally, for the Bubble rule, the scaling coefficient estimated was 0.74. Naively, one would think that the sorting mechanism of the Bubble rule would take on average 2/3*L* time steps, because that would be the average time for the rightmost blue state or the leftmost red state to move to its respective boundary position. However, because of the resolution module needed to solve the ambiguities of sorting, this process is delayed. While the red states in the resolution module just keep moving to the right as a black state, the blue state contained in the black state is delayed by exactly 1 time step, since it leaves the black state vertically instead of diagonally (fig. 4d). To understand how long the pattern formation process takes, then, we need to estimate into how many resolution modules the leftmost blue state enters during its sorting. First, whether or not a blue and a red state with only whites or blues in between need to be resolved is a matter of parity. If the states are at an even distance, they will clash and need to be resolved; if they are at an odd distance, they will just be swapped when directly neighboring and do not need the resolution module. Now, as the red states are propagating to the right, any initially neighboring red states need to create a gap between them in order to continue their sorting. Therefore, red states being direct neighbors or next nearest neighbors will be in the same parity, preventing the second one from needing to use a resolution module no matter where the leftmost blue state starts. Taking that into account, the number of resolution modules into which the rightmost blue state enters can be estimated as the number of red states it meets (on average *L/*3) times the probability that that red state is at least 3 positions away from the previous one ((2/3)^2^) times the probability that that red state is in the right parity (1/2). Overall, then, the number of resolution modules is on average 2*L/*27 and the mean time to steady state for this rule is 2*L/*3 + 2*L/*27 = 20*L/*27 ≈ 0.74*L*, as found numerically (cf. fig. S1).

**Appendix 5 - supplementary figure S1.**
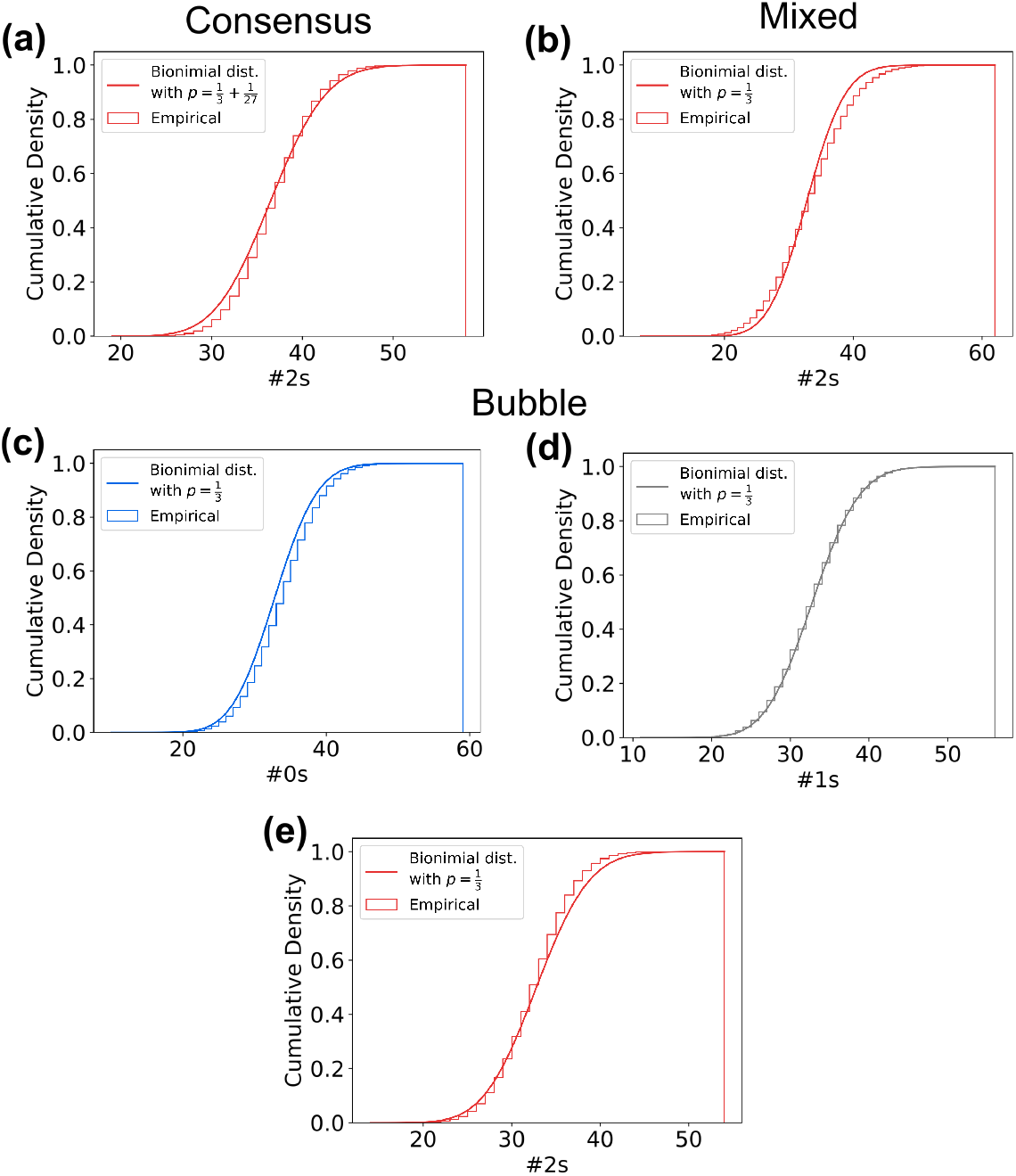
Distribution of different states after the 2*L* steps for *L* = 99 of the Consensus (a), Mixed (b) and Bubble (c-e) rule. Empirical distributions where sampled from 100000 samples, binomial distributions plotted with indicated probability *p* (see Analytical models). Although in some cases the binomial appears to be extremely close to the empirical distribution, the Kolmogorov-Smirnov-Tests fail in all cases (*p* << 0.001).

## Notes

### Competing Interest Statement

The authors have declared no competing interest.

